# PHF19 drives PRC2 sub-nuclear compartmentalization to promote motility in TNBC cells

**DOI:** 10.1101/2025.03.13.642950

**Authors:** Nina Pelzer, Teodora Lukic, Wanwan Ge, Nina Schnabel, Peter Teufel, Monilola A. Olayioye, Zeynab Najafova, Cristiana Lungu

## Abstract

Polycomb Repressive Complex 2 (PRC2) is a key regulator of chromatin architecture and transcriptional repression, playing essential roles in development and disease. While its enzymatic activity is well characterized, the molecular factors governing PRC2 subnuclear organization, particularly in cancer cells, are largely unknown. Here, we integrate high-resolution *in situ* spatial proteomics, imaging, and functional genomics to investigate PRC2 compartmentalization in triple-negative breast cancer (TNBC) cells. We identify PHF19, a sub-stoichiometric PRC2 accessory subunit upregulated in TNBC, as a key factor driving the formation of micron-sized nuclear PRC2 bodies. These structures act as functional hubs that stabilize PRC2 occupancy and reinforce H3K27me3 domain organization. Mechanistically, we identify an intrinsically disordered region (IDR) within PHF19 that is essential for its clustering behavior in cells and link this property to the role of PHF19 in promoting cancer cell motility. Our findings uncover a non-enzymatic layer of PRC2 regulation, whereby its local compartmentalization through accessory subunits directly impinges on cellular behavior. These insights expand our understanding of PRC2 spatial biology with implications for both normal development and disease progression.

## Introduction

Polycomb group (PcG) proteins are master regulators of gene expression and chromatin organization, playing critical roles in development, cellular differentiation, and disease ^1^. These proteins are primarily organized into two major complexes: Polycomb Repressive Complex 1 (PRC1) and PRC2, both of which mediate transcriptional repression of target genes. PcG proteins achieve this repression by modifying chromatin architecture, either through histone modifications or higher-order chromatin organization ^1,2^.

Through its catalytic subunit Enhancer of Zeste Homolog 2 (EZH2), PRC2 catalyzes the methylation of histone H3 at lysine 27 (H3K27me1/2/3), a hallmark of repressive chromatin that serves as a platform for downstream silencing mechanisms ^3^. In mammals, PRC2 assembles into two mutually exclusive protein subcomplexes, PRC2.1 and PRC2.2. PRC2.1 contains one of three Polycomb-like proteins (PHF1, MTF2, or PHF19) along with either PALI1/2 or EPOP ^4–8^. PRC2.2, in contrast, incorporates JARID2 and AEBP2 ^8–10^. Although both PRC2.1 and PRC2.2 contribute to H3K27me3 deposition, they adopt distinct genomic targeting strategies that collectively shape the H3K27me3 landscape and guide canonical PRC1 (cPRC1) recruitment ^11^. Specifically, PRC2.1 binds in sharp peaks at CpG islands and catalyzes the majority of H3K27me3 at Polycomb target genes. Conversely, PRC2.2 is poor at catalyzing H3K27me3, it mirrors the H2AK119ub1 distribution, and is essential for the recruitment of CBX7-cPRC1^11^.

While the enzymatic activities of PcG proteins have been extensively studied, increasing evidence highlights the importance of their subcellular organization as an additional layer of functional regulation ^2,12–14^. For example, in embryonic stem cells (ESCs), PcG proteins accumulate in optically resolved nuclear bodies, commonly referred to as Polycomb bodies. These specialized nuclear compartments concentrate PcG activity at specific target loci, such as developmentally regulated genes ^12,15–19^. Polycomb bodies are thought to be important for maintaining chromatin organization, gene repression memory, and play a role in developmental state transitions ^20–22^. Intriguingly, these structures are dynamic and lose cohesiveness during differentiation ^12,15–17,21,23,24^, yet the molecular mechanisms driving their assembly, maintenance, and dissolution, are not fully understood. Additionally, some PcG components also exhibit noncanonical functions in the cytoplasm. For example, cytosolic EZH2 has been shown to regulate actin polymerization in immune cells and fibroblasts ^25,26^, underscoring the intimate connection between PcG spatial organization and cellular context.

Importantly, the molecular mechanisms that regulate the subcellular localization of PRC2 subunits can be hijacked in different disease settings. Specifically, in triple-negative breast cancer (TNBC), a highly metastatic BC subtype, EZH2 was reported to be phosphorylated at Thr367, leading to its cytoplasmic export and the activation of signaling pathways that enhance metastatic dissemination ^27–29^. While cytosolic mislocalization of EZH2 in cancer cells is well-documented, it remains unclear whether and how EZH2 complexes may be reorganized at the subnuclear level in such diseased cells. Addressing this question requires approaches capable of unbiased profiling of endogenous epigenetic compartments, with high spatial resolution and without interference with the native cellular balance.

Here, we address this knowledge gap in Polycomb biology by investigating the molecular mechanisms underlying PRC2 subnuclear organization in TNBC model systems. Leveraging unbiased *in situ* spatial proteomics, we identified PHF19, a sub-stoichiometric subunit of PRC2, as a key factor driving the formation of micron-sized PRC2 nuclear bodies in TNBC cells. By combining cellular perturbation experiments, high-resolution imaging, and epigenomic profiling, we demonstrate that PHF19 is not only essential for the subnuclear clustering of PRC2, but also coordinates the spatial organization of H3K27me3 domains in these cells. Notably, this function is unique to PHF19 and cannot be compensated for by the other PCL family members, MTF2 and PHF1. Mechanistically, we identify a previously undescribed intrinsically disordered region (IDR) within PHF19 that facilitates its compartmentalization into nuclear bodies. Functionally, we find that this spatial organization is important for the role of PHF19 in promoting TNBC cell motility.

Together, our findings reveal a novel role for PHF19 in shaping PRC2 nuclear organization and highlight subnuclear compartmentalization through accessory subunits as a key regulatory mechanism in Polycomb biology, with implications for epigenetic regulation in both normal development and disease contexts.

## Results

### Endogenous PRC2 core subunits accumulate in micron-sized nuclear bodies in bone entrained TNBC cells

To identify novel regulatory mechanisms underlying the spatial organization of PRC2 in cancer cells, we utilized MDA-MB-231 cells, a well-established TNBC model system previously employed to study EZH2 cytoplasmic localization ^29–31^. This cell line offers the additional advantage of available isogenic sub-populations with organ-specific metastasis preferences, providing a tractable model to explore PRC2 spatial regulation in distinct cellular contexts. Specifically, we used the BoM-1833 cell line, which was derived by intracardiac injection of parental MDA-MB-231 cells into mice, followed by isolation of a sub-population that preferentially metastasizes to the bone niche ^32^.

We first employed high-resolution microscopy to analyze the endogenous localization of PRC2 core subunits in the two cellular models. At a lateral resolution of 50 nm, both EZH2 and SUZ12 showed a diffuse distribution within the nuclei of MDA-MB-231 cells. In contrast, BoM-1833 cells displayed a striking accumulation of both PRC2 subunits within discrete nuclear bodies of approximately 1 µm in diameter (Figure 1A-C). These structures, resembling Polycomb bodies previously described in ESCs ^33^ and thereafter referred to as “PRC2 bodies”, were observed in approximately 50% of the BoM-1833 cells (Figure 1D) and overlapped with H3K27me3 foci (Figure 1E). These findings suggest that the PRC2 bodies we identified in BoM-1833 cells are not dysfunctional protein aggregates but rather reflect cell context-dependent functions of PRC2.

**Figure 1:**
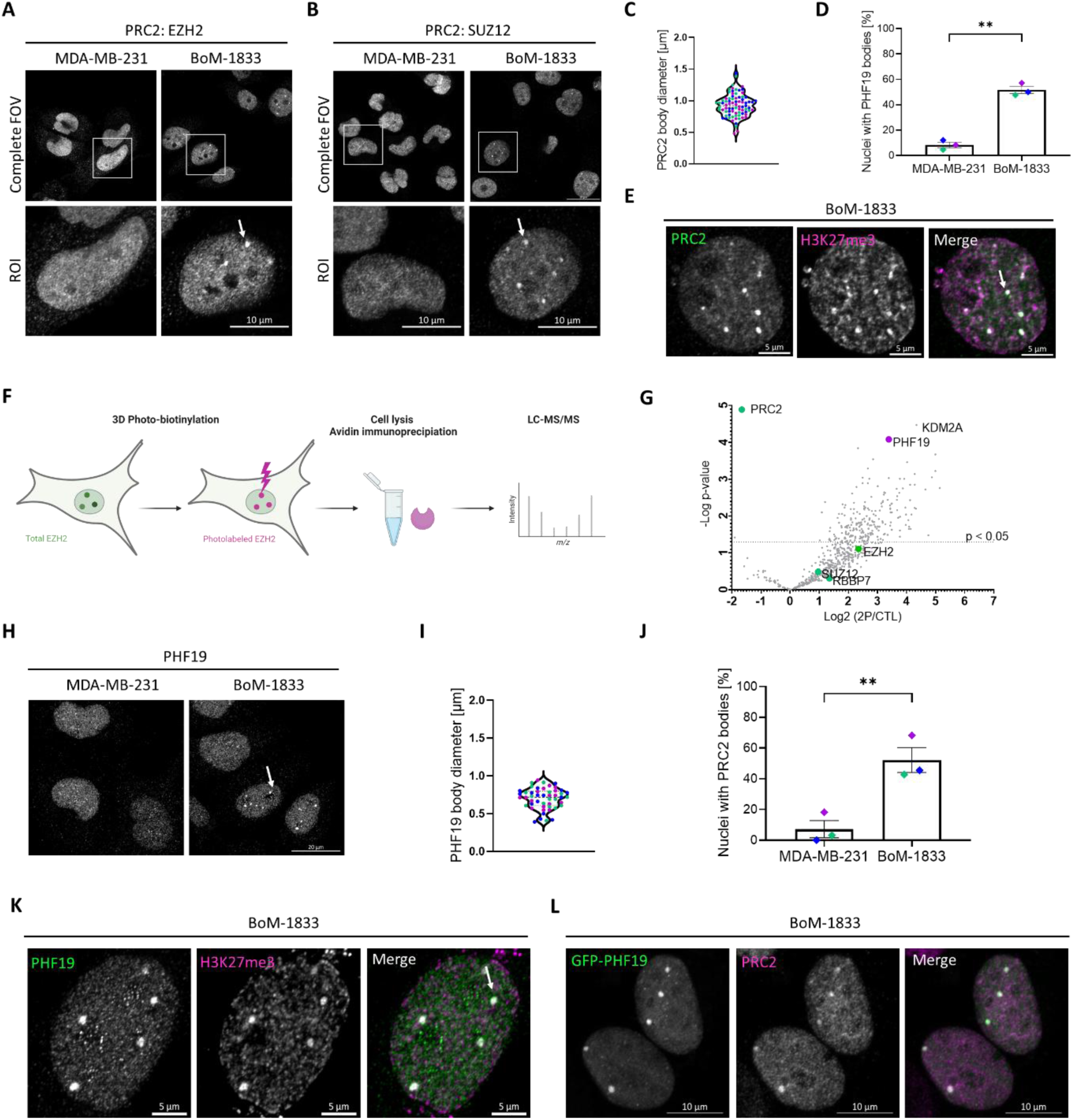
PRC2 core subunits accumulate within PHF19-positive nuclear bodies in bone entrained TNBC cells. (**A-B**) Representative confocal fluorescence microscopy images of endogenous EZH2 (A) or SUZ12 (B) immunostaining in MDA-MB-231 and BoM-1833 cells. Insets highlight exemplary nuclear bodies of EZH2 or SUZ12 accumulation (arrows) in the BoM-1833 cells. Scale bar: 10 µm. Images were acquired and are displayed with identical settings. (**C**) Violin plot quantifying PRC2 body diameter in BoM-1833 cells. Each dot represents a single PRC2 body; data from 3 biological replicates (N = 16–32 cells). (**D**) Quantification of percentage of cell nuclei with PRC2 bodies in MDA-MB-231 and BoM-1833 cells, based on the images representatively shown in A-B. Data represent measurements from n = 3 biological replicates. Biological repeats are color coded. Statistical significance was determined via unpaired t-test, p=0.0102. Error bars indicate mean ±SEM. (**E**) Representative confocal fluorescence microscopy image of BoM-833 cells stained for endogenous PRC2 (SUZ12, green) and H3K27me3 (magenta) immunostaining in BoM-1833 cells. The arrow indicates an exemplary area of co-localization at a PRC2 body. Scale bar: 5 µm. (**F**) Schematic representation of the 3D photo-biotinylation approach used to map the proteome of endogenous PRC2 bodies. Total EZH2 (green) is spatially distributed within the cell and selectively photo-biotinylated at defined regions of interest (magenta) upon light activation. Following cell lysis, biotinylated proteins are captured using avidin-based immunoprecipitation and analyzed by liquid chromatography-tandem mass spectrometry (LC-MS/MS). The figure was created using Biorender. (**G**) Volcano plot illustrating the proteomic content of PRC2 bodies in BoM-1833 cells. Analysis was performed on the 1384 proteins identified as enriched in the labeled versus control condition in all 4 biological repeats, with unique peptides ≥ 2, fold change ≥ 1.5; and t-test significance ≤ 0.05. The x-axis represents the log_2_ enrichment ratio (2P/CTL), and the y-axis represents the -log_10_ p-value, indicating statistical significance. The dotted horizontal line corresponds to the p-value threshold (p < 0.05). Members of the core PRC2 complex are labeled in green. (**H**) Representative confocal fluorescence microscopy images of endogenous PHF19 immunostaining in MDA-MB-231 and BoM-1833 cells. The arrow highlights exemplary accumulations of PHF19 within nuclear bodies in BoM-1833 cells. Scale bar: 20 µm. The images were acquired and are displayed with identical settings. (**I**) Violin plot showing the quantification of endogenous PHF19 body diameter in BoM-1833 cells based on the images representatively shown in (H). Data represent measurements from N = 14–17 cells across n = 3 biological replicates, with each dot representing the diameter of a single PHF19 body. Biological repeats are color coded. (**J**) Quantification of percentage of cell nuclei with PHF19 bodies in MDA-MB-231 and BoM-1833 cells, based on the images representatively shown in (I). Data represent measurements from n = 3 biological replicates. Biological repeats are color coded. Statistical significance was determined via unpaired t-test, p=0.003. Error bars indicate mean ±SEM. (**K**) Representative confocal fluorescence microscopy image of endogenous PHF19 (green) and H3K27me3 (magenta) immunostaining in BoM-1833 cells. The arrow indicates an exemplary area of co-localization at a PHF19 body. Scale bar: 5 µm. (**L**) Representative confocal fluorescence microscopy images of BoM-1833 cells, 24 h post transfection with a GFP-PHF19 (green) expression plasmid and immunostained for endogenous core PRC2 subunits (SUZ12, purple). The arrow indicates an exemplary area of co-localization. Scale bar: 10 µm.

### Unbiased spatial proteomics identifies PHF19 as an integral part of the PRC2 bodies

We hypothesized that the nuclear bodies we observe in BoM-1833 cells, might contain regulatory proteins that govern the distinct subcellular organization of PRC2. To address this, we employed opto-proteomics ^34,35^, a cutting-edge spatial proteomics method that integrates microscopy-guided protein labeling with mass spectrometry. This technology enables the unbiased identification of proteins localized to optically resolved cellular compartments without the need of genetic engineering (Figure 1F). Using EZH2 staining to identify endogenous PRC2 bodies (Figure S1A), we applied targeted 3D photo-biotinylation to selectively label proteins within these nuclear structures, *in situ*. This approach enhances spatial specificity while minimizing background noise, enabling precise proteome identification with a resolution of 250 nm ^34,35^ (Figure S1B-C).

1,384 unique protein hits were detected as common across the four biological replicates, marking a 56% overlap (Figure S1D and Table S1). REACTOME pathway analysis of these hits, filtered using stringent criteria (Figure S1E and Materials and Methods), revealed an enrichment of proteins involved in chromatin-related processes, including PRC2-mediated histone methylation. Notably, core PRC2 subunits EZH2 and SUZ12 as well as RBBP7 were consistently enriched in the illuminated (2P) samples compared to non-illuminated (CTL) controls across all four biological replicates (Figure 1G). These findings showcase the high reproducibility and specificity of the opto-proteomics approach in defining the molecular composition of the PRC2 bodies.

Further ranking of the proteome data according to the significance of peptide enrichment in the biotinylated condition relative to the unlabeled control, identified KDM2A and PHF19 as the most enriched candidates (Figure 1G). KDM2A plays a key role in regulating chromatin accessibility and gene expression by removing H3K36me2, a mark that interferes with the recruitment of PRC2 to chromatin ^36,37^. PHF19, a sub-stoichiometric subunit of the PRC2.1 complex, belongs to the PCL family, which also includes MTF2 and PHF1. It recognizes H3K36me2/3 marks and facilitates PRC2 recruitment to specific genomic loci, thereby enhancing H3K27me3 deposition and transcriptional repression ^38–41^. While PHF19 is a known regulator of PRC2 activity ^3^, its molecular function in breast cancer cells has not been explored so far. To address this gap, we thereby focused on investigating its functional relevance in this context.

Immunostaining of endogenous PHF19, using a knockdown-validated antibody (Figure S2A), revealed a predominantly diffuse localization pattern of the protein in the nuclei of parental MDA-MB-231 cells (Figure 1H). By contrast, in the BoM-1833 sub-population, around 40% of the cells displayed an accumulation of PHF19 in distinct nuclear bodies of approximately 0.8 µm in diameter (Figure 1H-J). These structures overlapped with H3K27me3 foci (Figure 1K). Further analysis demonstrated that GFP-tagged PHF19 also formed nuclear bodies in BoM-1833 cells, which co-localized with endogenous PRC2 core subunits (Figure 1L). Together, these findings suggest that PHF19 may play a role in mediating the context-specific PRC2 regulation in TNBC cells.

### PHF19 controls the subcellular organization of PRC2 bodies in bone entrained TNBC cells

To investigate whether PHF19 is important for the structural organization of PRC2 bodies in BoM-1833 cells, we performed transient PHF19 depletion experiments using two independent siRNAs. Strikingly, PHF19 knockdown was paralleled by a dramatic loss of PRC2 nuclear bodies, as revealed by immunofluorescence staining (Figure 2A-B). Quantitative analysis showed that the percentage of cells exhibiting PRC2 foci dropped to under 1% upon PHF19 depletion (Figure 2C). Importantly, the expression levels of core PRC2 subunits EZH2 and SUZ12 were unaffected, indicating that the PRC2 complex remains intact under these conditions (Figure 2D-G). Furthermore, while the other two PCL family members, PHF1 and MTF2, were expressed in BoM-1833 cells, their levels were not altered by PHF19 knockdown (Figure 2H-I). These findings demonstrate that PHF19 is critical for maintaining the spatial organization of PRC2 bodies in BoM-1833 cells. Furthermore, this function appears to be unique to PHF19, as it cannot be compensated for by the other PCL family members.

**Figure 2:**
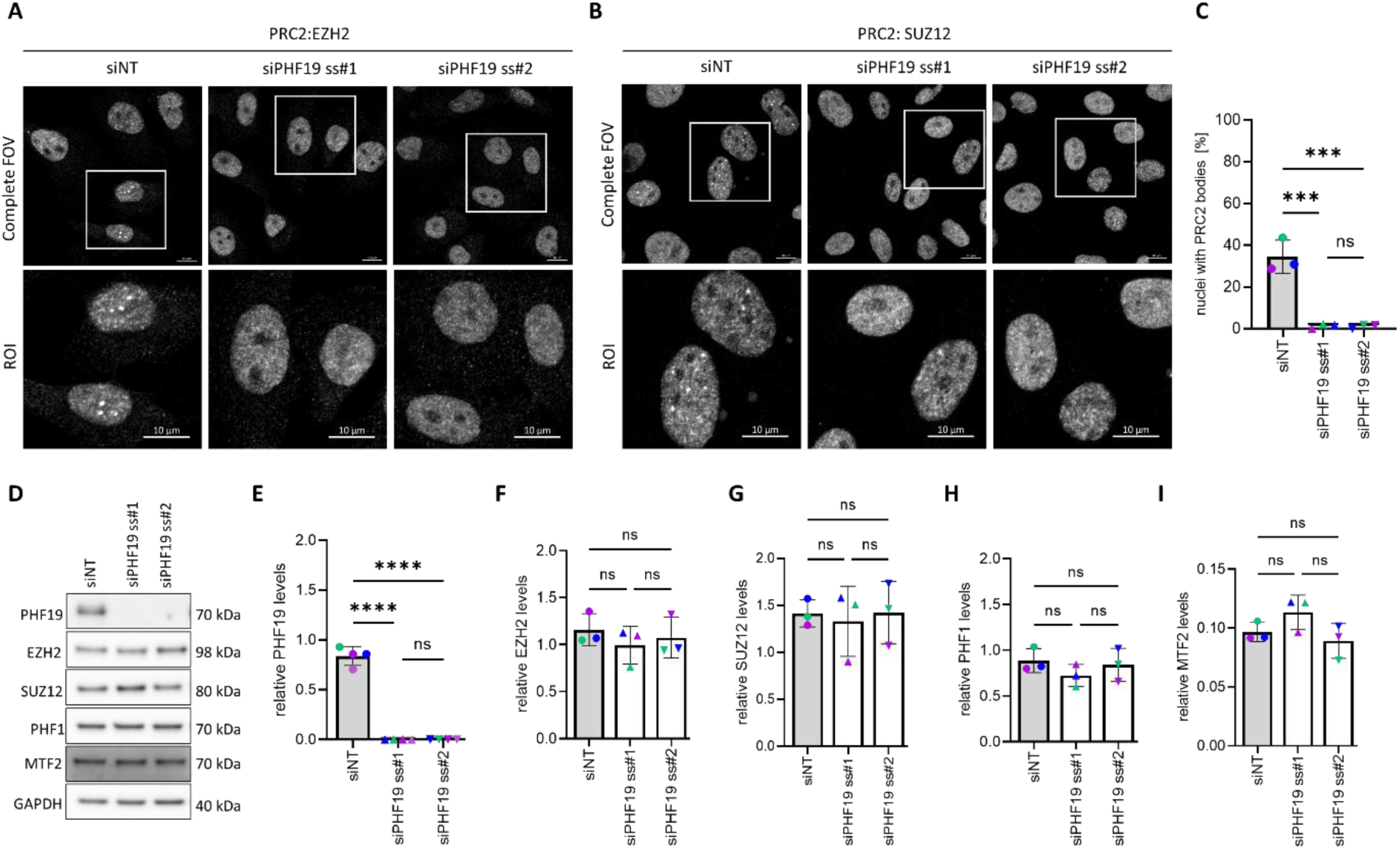
PHF19 controls the organization of PRC2 bodies in bone entrained TNBC cells. (**A-B**) Representative confocal fluorescence microscopy images of BoM-1833 cells transfected with the indicated siRNAs. Cells were fixed 96 hours post-transfection and immunostained for endogenous EZH2 (A) or SUZ12 (B). Regions of interest (ROIs) are highlighted, with inset images showing magnified views of the immunostained cells. Scale bar: 10 µm. Images that are to be directly compared where imaged and are displayed with identical settings. (**C**) Quantification of the percentage of nuclei exhibiting PRC2 bodies in BoM-1833 cells treated as in (A-B) and immunostained for PRC2 core subunits. Data represent measurements from N = 50–60 cells across n = 3 biological replicates. Biological repeats are color coded. Statistical significance was determined via one-way ANOVA testing, *** = 0.0003, ns= not significant. Error bars indicate mean ±SD. (**D**) BoM-1833 cells were transfected with the indicated siRNAs and lysed 96 hours later for Western blot analysis using the specified antibodies. GAPDH was used as loading control. (**E-I**) Densitometric analysis of PHF19 (E), EZH2 (F), SUZ12 (G), PHF1 (H) and MTF2 (I) protein levels in cell lysates obtained from BoM-1833 cells treated as described in (D). GAPDH was used for relative normalization of the chemiluminescence signal obtained for the different PRC2 subunits. Data represent measurements from n = 3 biological replicates, whereby the values for siPHF19 are reported relative to the mean value of the control (siNT) within each biological replicate. Biological repeats are color coded. Statistical significance was determined via one-way ANOVA testing, **** < 0.0001, ns = not significant. Error bars indicate mean ±SD.

### PHF19 is sufficient to drive the subcellular organization of PRC2 bodies in TNBC cells

We next investigated whether the role of PHF19 in regulating PRC2 subnuclear compartmentalization extends to other cell systems. Analysis of a TCGA breast cancer dataset, stratified by molecular subtype, revealed elevated *PHF19* expression in most tumor samples, with a statistically significant upregulation in TNBC patients (Figure 3A). In contrast, *PHF1* was downregulated in BC, while *MTF2* showed no significant expression changes (Figure S3).

**Figure 3:**
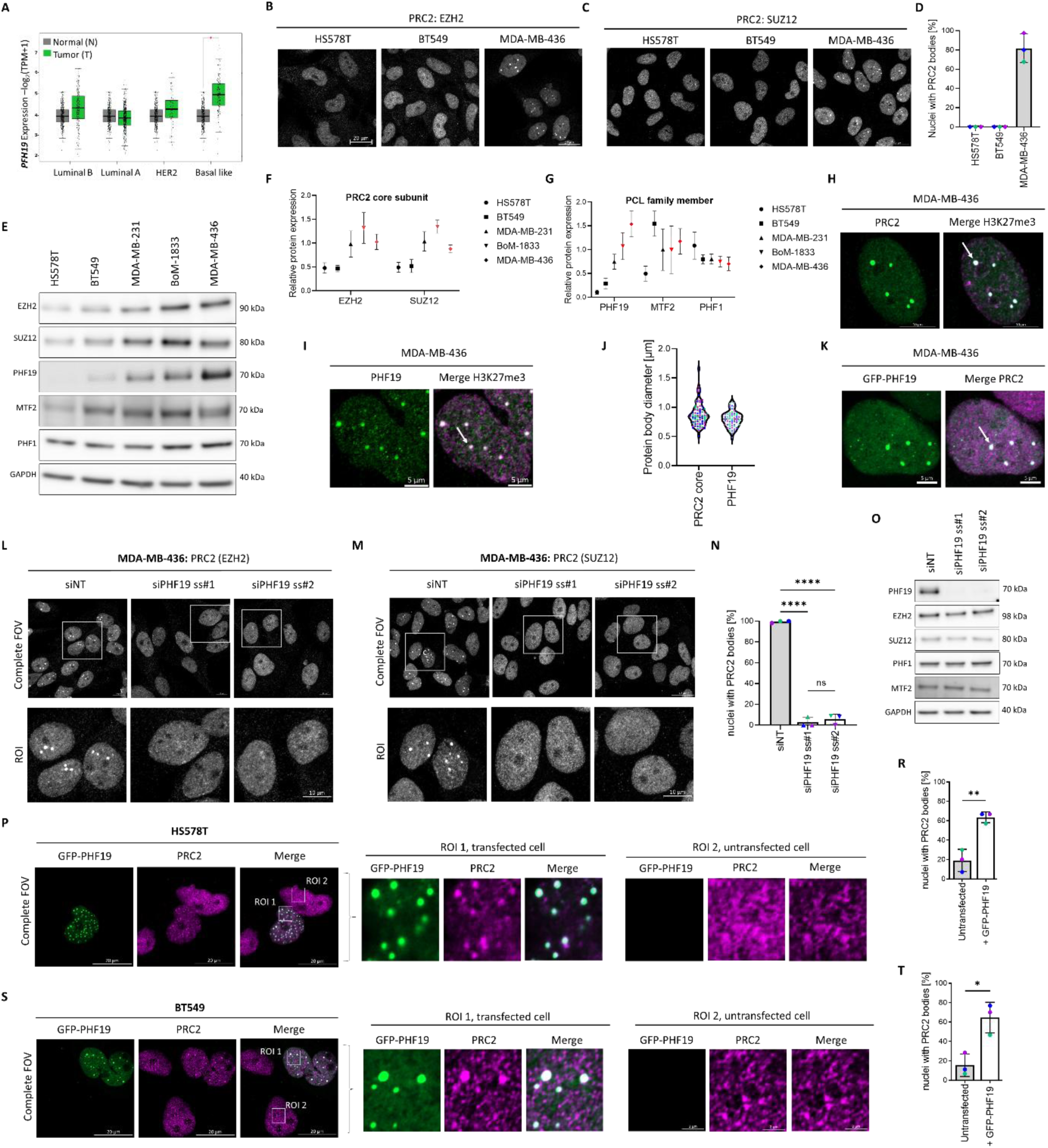
PHF19 controls the organization of PRC2 bodies in TNBC cells. (**A**) *PHF19* gene expression analysis across a TCGA BRCA cohort sorted by molecular subtype subtype. Box plots display the expression levels of *PHF19* in normal (grey) and tumor (green) tissue for the indicated breast cancer subtypes. Data are derived from TCGA/GTEx datasets and visualized using GEPIA2. Statistical significance between tumor and normal samples was determined by unpaired t-test (*p < 0.05). n= 291 (Normal), 194 (Luminal B), 415 (Luminal A), 66 (HER2), 135 (Basal-like). (**B-C**) Representative confocal microscopy images of EZH2 (B) and SUZ12 (C) immunostaining in the indicated cell lines. Scale bar: 20 µm. Images that are to be directly compared were recorded and are displayed using identical settings. (**D**) Quantification of the percentage of cell nuclei with PRC2 bodies in the indicated cell lines based on confocal microscopy images as shown in (B-C). Data represent measurements from N = 35– 55 cells across n = 3 biological replicates. Biological repeats are color coded. (**E**) Representative immunoblot analysis of full cell lysates prepared from the indicated cell lines and using the annotated antibodies. GAPDH was used as the loading control. (**F-G**) Densitometric quantification of EZH2, SUZ12 (F) and PCL family (G) subunit protein expression in the TNBC cell line panel used in this work. GAPDH was used for normalization of the chemiluminescence signal of the PRC2 subunits across cell lines. The data for siPHF19 are reported relative to the mean values for the siNT control. Data represent measurements from n = 3 biological replicates, error bars are mean ±SD. Measurements stemming from cell lines forming detectable PRC2 bodies by Airyscan microscopy were highlighted in red. (**H-I**) Representative confocal fluorescence microscopy images showing co-immunostaining of H3K27me3 with the endogenous PRC2 core subunit SUZ12 (H) and PHF19 (I) in MDA-MB-436 cells. Arrows indicate exemplary regions of colocalization. Scale bar: 10 µm (H), 5 µm (I). (**J**) Violin plot showing the quantification of PRC2 core and PHF19 protein body diameter as based on the images representatively shown in (F-G). Data represent measurements from N = 14–29 (core PRC2 subunits) and N= 19-22 (PHF19) cells across n = 3 biological replicates, with each dot representing the diameter of a single protein body. Biological repeats are color coded. (**K**) Representative confocal fluorescence microscopy images of MDA-MB-436 cells, 24 h post transfection with GFP-PHF19 (green) and immunostained for endogenous SUZ12 (purple). The arrow indicates an exemplary area of co-localization. Scale bar: 5 µm. (**L-M**) MDA-MB-436 cells were transfected with the indicated siRNAs followed by fixation 96 h later and immunostaining for endogenous EZH2 (L) or SUZ12 (M). The bottom row shows magnified views of the cropped fields of view. Images that are to be directly compared were acquired and are displayed using identical settings. Scale bar: 10 µm (**N**) Quantification of percentage of cell nuclei with PRC2 bodies in MDA-MB-436 cells transfected with the indicated siRNAs and imaged as representatively shown in (L-M). Data represent measurements from n = 3 biological replicates. Biological repeats are color coded. Statistical significance was determined via one-way ANOVA, ****= 0.001, ns= not significant. Error bars indicate mean ±SD. (**O**) MDA-MB-436 were treated as described in (L-M), followed by cell lysis. The material was analyzed by Western blot using the indicated antibodies. See also Figure S4. (**P**, **S**) Representative confocal microscopy images and (**R**, **T**) quantification of HS578T (P, R) and BT549 (S, T) fixed 24 h after transfection with a plasmid encoding for GFP-PHF19 (magenta) and immunostained for endogenous SUZ12 (PRC2 core). ROIs (Regions of Interest) are highlighted and magnified, showing the endogenous localization of SUZ12 in cells transfected with GFP-PHF19 (ROI 1) versus un-transfected cells (ROI 2). Scale bar: 20 µm. The bar diagrams show the endogenous SUZ12 localization phenotype in relation to the GFP-PHF19 expression status. Data represent measurements from N = 7–30 cells from n = 3 biological replicates. Biological repeats are color coded. Statistical significance was determined via unpaired t-test, * = 0.0123, **= 0.0038. Error bars indicate mean ±SD.

Based on these findings, we focused on the TNBC cellular context to further examine the interplay between PHF19 and the subcellular compartmentalization of PRC2. We selected three additional cell models frequently used in TNBC research (HS578T, BT549, and MDA-MB-436) ^42–44^ and analyzed the localization of PRC2 core subunits EZH2 and SUZ12 (Figure 3B-C) by high resolution microscopy. Interestingly, while HS578T and BT549 cells exhibited a diffuse nuclear localization of both proteins, MDA-MB-436 cells displayed prominent PRC2 bodies in over 90% of the population (Figure 3B-D). Western blot analysis further revealed that the BoM-1833 and MDA-MB-436 cells express the highest levels of EZH2, SUZ12 and PHF19 (Figure 3E-G). By comparison, HS578T and BT549 cells, which do not show PRC2 bodies, have very low PHF19 protein levels. PHF1 and MTF2 were largely uniformly expressed across all cell lines (Figure 3E-G).

Given the presence of PRC2 bodies in MDA-MB-436 cells, we selected this cell line as a second model system to investigate the molecular function of PHF19 in PRC2 subcellular organization. Co-localization studies revealed that PRC2 bodies overlapped with H3K27me3 foci, indicating that these structures represent enzymatically active PRC2 complexes (Figure 3H). Additionally, endogenous PHF19 accumulated in discrete nuclear clusters that co-localized with H3K27me3 domains (Figure 3I and S2B). These foci, approximately 1 µm in diameter, were consistently observed for both endogenous PHF19 and PRC2 core subunits (Figure 3J). Moreover, GFP-tagged PHF19 accumulated at optically distinct compartments that overlapped with endogenous PRC2 bodies (Figure 3K), further supporting a regulatory cross-talk between PHF19 and PRC2 subnuclear organization in this model system. Indeed, consistent with the findings in the BoM-1833 cells, PHF19 depletion led to the complete dissolution of nuclear EZH2 and SUZ12 foci, with the percentage of MDA-MB-436 cells exhibiting PRC2 bodies dropping from nearly 100% to less than 0.1% (Figure 3L-N). Western blot analysis confirmed efficient knockdown of PHF19 without affecting the expression of core PRC2 subunits (EZH2 and SUZ12) or other PCL family members (PHF1 and MTF2) (Figure 3O, Figure S4). These results demonstrate that PHF19 is essential for maintaining the structural integrity of PRC2 bodies in both BoM-1833 and MDA-MB-436 cells, a function that cannot be compensated for by PHF1 and MTF2.

We next tested whether PHF19 is sufficient to drive PRC2 clustering *de novo*. To this end, we transfected GFP-tagged PHF19 into the low-PHF19-expressing cell lines HS578T and BT549, where endogenous PRC2 core subunits exhibit a largely diffuse nuclear localization pattern at the resolution of our assay. In both models, GFP-PHF19 was able to form multiple, prominent nuclear clusters. These overlapped with newly formed PRC2 bodies (Figure 3P & S). Notably, approximately 60% of the GFP-PHF19 transfected cells displayed PRC2 bodies (Figure 3R & T), representing a threefold increase compared to un-transfected cells. Together, our findings establish PHF19 as a central driver of PRC2 subnuclear organization in TNBC cells.

### PHF19 clustering at PRC2 target sites requires the cooperation of the IDR, the DNA binding and the SUZ12 interaction domains

We next sought to elucidate the biochemical basis underlying the clustering behavior of PHF19. To this end, we performed imaging experiments with MDA-MB-436 cells, since they display prominent endogenous clusters for both PHF19 and the core subunits of PRC2 (Figure 3). Notably, upon transient transfection of GFP-PHF19, we observed that the expression levels of the protein directly influenced its subcellular localization (Figure 4A-B). Specifically, at low expression levels, GFP-PHF19 formed a small number of prominent clusters that largely overlapped with H3K27me3 foci. By contrast, at high expression levels, numerous smaller clusters were visible, also outside of the H3K27me3 foci. This phenotype suggests saturation of the endogenous core PRC2 machinery, likely reflecting the presence of both homotypic PHF19-only compartments and heterotypic PHF19-PRC2 clusters.

**Figure 4:**
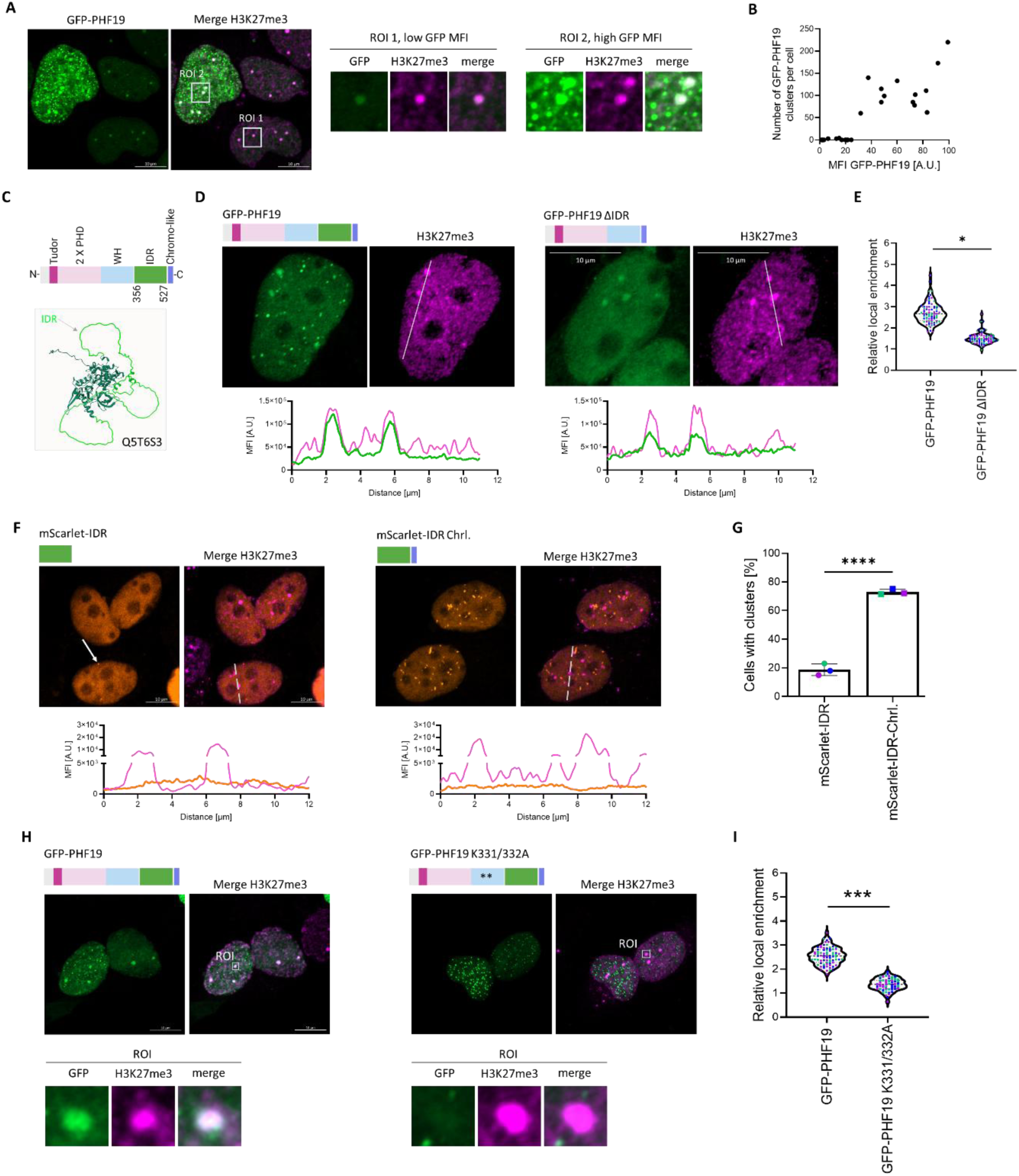
PHF19 clustering at PRC2 targets requires the cooperation of the IDR, the WH and the Chromo-like domain. (**A**) Representative confocal microscopy image of MDA-MB-436 cells fixed 24 hours after transfection with GFP-PHF19 (green) and co-immunostained for H3K27me3 (magenta). Insets show zoomed-in view of two regions of interest (ROI), selected based on the GFP-PHF19 expression levels of the corresponding transfected cells. Scale bar: 10 µm. (**B**) Dot blot showing the relationship between GFP-PHF19 mean fluorescence intensity (MFI) and the number of GFP-PHF19 clusters per cell. The quantification is based on the confocal microscopy images representatively shown in (A). Each dot represents a single cell nucleus. N= 26 (**C**) Top – Schematic representation of the PHF19 protein domain structure. PHD - plant homeodomain; WH - winged helix domain; IDR – intrinsically disordered region. Right – AlphaFold prediction of the PHF19 (Uniprot: Q5T6S3) protein structure, illustrating the presence of an IDR, highlighted in light green. (**D**) Representative confocal microscopy images of MDA-MB-436 cells fixed 24h after transfection with the indicated GFP-PHF19 variants and co-stained for H3K27me3. The histograms display the mean fluorescence intensity (MFI) across the line annotated in the merge channel. Scale bar: 20 µm. The images were acquired and are displayed with identical settings. (**E**) Quantification of the relative local enrichment of the indicated GFP-PHF19 variants at H3K27me3 foci, based on the images representatively shown in (D). The analysis reflects the normalized fluorescence intensity of GFP-PHF19 variants at endogenous PRC2 clusters relative to the surrounding nuclear background. Data represent measurements from N = 20– 29 cells across n = 3 biological replicates, where each dot represents the diameter of a single GFP-PHF19 body. Biological repeats are color coded. Statistical significance was determined using an unpaired t-test (p = 0.0199). (**F**) Representative confocal microscopy images of MDA-MB-436 cells fixed 24h after transfection with the indicated PHF19 variants and co-immunostained for H3K27me3. The arrow indicates a representative cell exhibiting a nuclear clustering phenotype of the mScarlet-IDR protein. The histograms display the mean fluorescence intensity (MFI) across the line annotated in the merge channel. Scale bar: 10 µm. (**G**) Quantification of the PHF19 domains localization phenotype in MDA-MB-436 cells transfected with the specified protein variants. The bar graph depicts the percentage of cells exhibiting a clustered phenotype, as observed in the confocal microscopy images shown in (F). Data represent measurements from N = 35–52 cells across n = 3 biological replicates. Biological repeats are color coded. Statistical significance was determined using an upaired t-test (p <0.0001). (**H**) Representative confocal microscopy images of MDA-MB-436 cells fixed 24h after transfection with the indicated PHF19 variants and co-immunostained for H3K27me3. Scale bar: 10 µm. Insets show a zoomed-in view of the regions of interest (ROI). (**I**) Quantification of the relative local enrichment of the indicated GFP-PHF19 variants, based on the images representatively shown in (H). Analyzed was the mean GFP fluorescence intensity at H3K27me3 bodies relative to the surrounding nuclear background. Data represent measurements from N = 19–20 cells across n = 3 biological replicates, with each dot representing the diameter of a single protein body. Biological repeats are color coded. Statistical significance was determined using an unpaired t-test (p = 0.0002).

The observation that the PHF19 clustering behavior is modulated by protein expression levels is reminiscent of liquid-liquid phase separation (LLPS) driven processes. This phenomenon that has been previously described for various Polycomb group proteins, including PHF1 ^14,45–49^. However, the specific molecular determinants driving PHF19 clustering, have not been studied so far. To address this, we analyzed the structural properties of PHF19 using AlphaFold, which predicted a long intrinsically disordered region (IDR) located between the DNA-binding and chromo-like domains (Figure 4C). Since IDRs are known to drive biomolecular condensate formation and process specificity ^50^, we hypothesized that IDR of PHF19 could play a role in its localization to nuclear bodies.

To test this, we generated a PHF19 mutant lacking the IDR (ΔIDR) and compared its localization to the wild-type (WT) GFP-PHF19 protein. At similar expression levels, the ΔIDR mutant exhibited a more diffuse nuclear distribution by comparison to WT PHF19. Nevertheless, a weak enrichment at H3K27me3 foci was still observable in MDA-MB-436 cells (Figure 4D). Quantitative analysis showed that WT PHF19 was enriched approximately 3-fold over background levels at H3K27me3 foci, while the ΔIDR mutant exhibited only 1.5-fold enrichment (Figure 4E). These findings indicate that the IDR facilitates robust accumulation of PHF19 at H3K27me3 domains, though it is not the sole determinant of this localization.

Next, we tested whether the IDR alone is sufficient for PHF19 clustering in cells. When expressed as a fusion with mScarlet, the IDR formed foci but only in a small fraction of cells (Figure 4F-G). However, extending the construct to include the chromo-like domain, which interacts with SUZ12 ^3,51^, drastically increased the frequency of clustering, with approximately 80% of transfected cells displaying foci (Figure 4F-G). Notably, these clusters did not overlap with H3K27me3 foci, suggesting that additional domains are required for targeting PHF19 to genomic loci pre-marked by active PRC2.

PCL proteins contain DNA-binding domains that exhibit some sequence preference and aid in the recruitment of PRC2 to chromatin ^40,41,52,53^. To investigate the role of DNA binding in PHF19-mediated clustering, we generated a loss-of-function mutant (K331/K332A) targeting the DNA-binding domain. Although this mutant retained the ability to form nuclear bodies, its enrichment at H3K27me3 sites was only slightly above background levels (Figure 4H-I).

Together, these findings indicate that PHF19 clustering is mediated by multiple structural features, with the DNA-binding domain facilitating targeting to specific genomic regions and the IDR promoting the local protein accumulation on these sites. These mechanisms likely function in a positive feedback loop with H3K27me3 and other histone marks such as H3K36me2/me3 to reinforce PRC2 clustering and activity.

### PHF19 controls the spatial organization of H3K27me3 domains without affecting global PRC2 activity in TNBC cells

To determine whether PHF19 influences not only PRC2 compartmentalization but also its enzymatic activity, we examined global and local H3K27me3 levels in BoM-1833 and MDA-MB-436 cells following PHF19 depletion. Western blot analysis revealed that PHF19 knockdown over four days had no significant effect on total H3K27me3 levels in either cell line (Figure 5A-B). In contrast, EZH2 depletion over the same period caused a pronounced reduction in H3K27me3 levels (Figure S5A-B). Remarkably, immunofluorescence analysis revealed that transient PHF19 knockdown led to the dissolution of the large H3K27me3 foci present in control cells (Figure 5C & E). In BoM-1833 cells, the percentage of cells displaying H3K27me3 clusters larger than 0.5 µm dropped from ∼40% in control conditions to <2% following PHF19 depletion (Figure 5D). Similarly, in MDA-MB-436 cells, PHF19 depletion caused a marked reduction in the number of H3K27me3 foci larger than 1 µm (Figure 5E-F). Further quantitative image analysis of MDA-MB-436 cells, which have more optically defined H3K27me3 foci, revealed a fragmentation towards smaller, more dispersed H3K27me3 clusters following PHF19 depletion (Figure 5F). Together, these results indicate that PHF19 is critical for maintaining the spatial organization of H3K27me3 domains.

**Figure 5:**
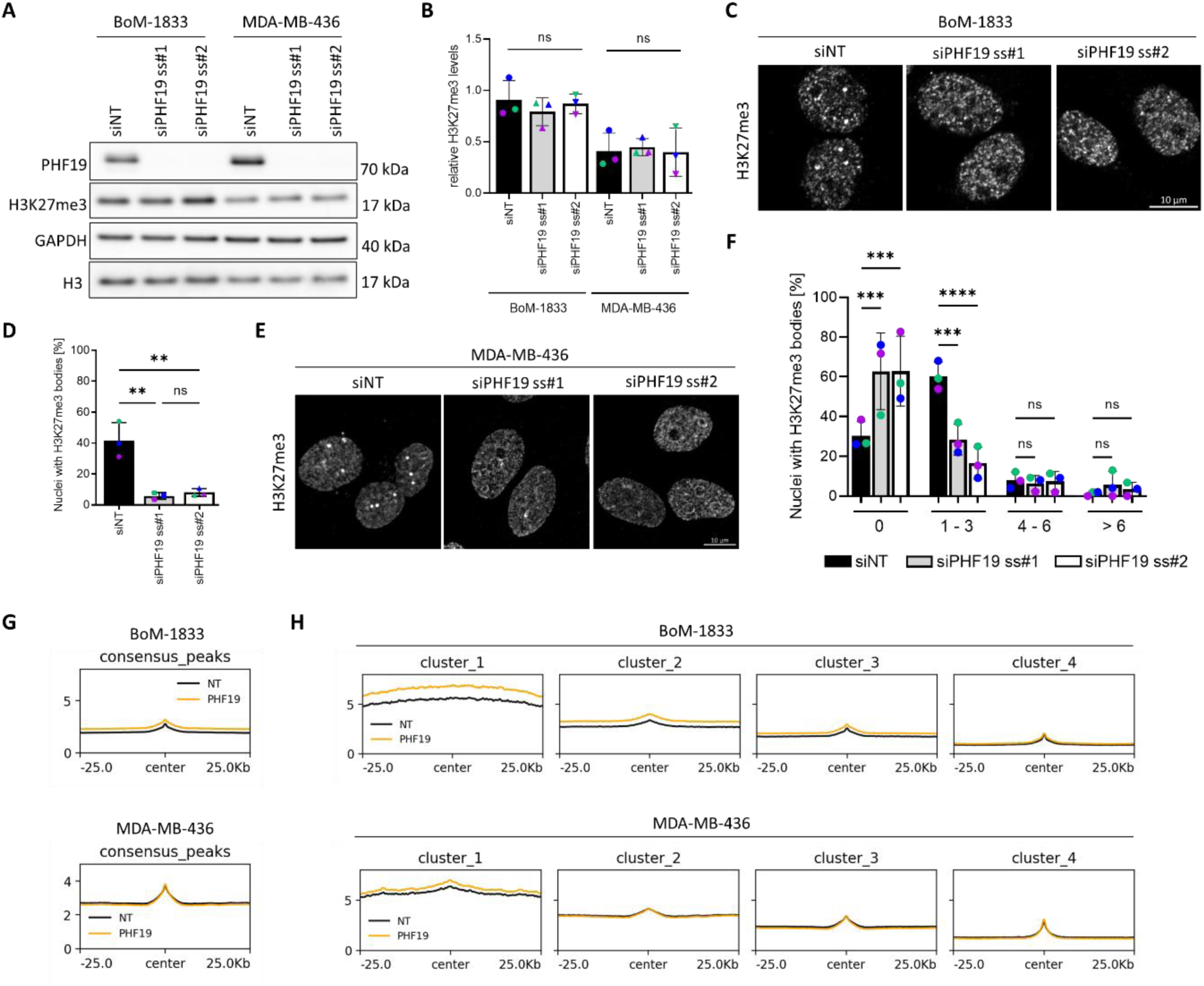
PHF19 controls the subcellular organization of H3K27me3 domains in TNBC cells. (**A**) BoM-1833 and MDA-MB-436 cells were transfected with the indicated siRNAs and lysed 96 hours later for Western blot analysis using the specified antibodies. GAPDH and H3 were used as loading controls. (**B**) Densitometric analysis of H3K27me3 protein levels in cell lysates obtained from BoM-1833 and MDA-MB-436 cells treated as described in (A). Data represent measurements from n = 3 biological replicates. Biological repeats are color coded. Statistical significance was determined via one-way ANOVA, ns= not significant. Error bars indicate mean ±SD. (**C, E**) Representative confocal microscopy images of BoM-1833 (C) and MDA-MB-436 (E) cells fixed 96h after transfection with the indicated siRNAs and immunostained for H3K27me3. Scale bar: 10 µm. Images that are to be directly compared were recorded and are displayed with identical settings. (**D**) Quantification of percentage of BoM-1833 cell nuclei with H3K27me3 bodies, based on the images representatively shown in (C). Data represent measurements from n = 3 biological replicates. Biological repeats are color coded. Statistical significance was determined via one-way ANOVA, **= 0.0022 (siNT vs siPHF19 ss#1); **= 0.0031 (siNT vs siPHF19 ss#2). ns= not significant. Error bars indicate mean ±SD. (**F**) Quantification of percentage of BoM-1833 cell nuclei with H3K27me3 bodies, based on the images representatively shown in E. Data represent measurements from n = 3 biological replicates. Biological repeats are color coded. Statistical significance was determined via two-way ANOVA **** < 0.0001, *** < 0.0008, ns= not significant. Error bars indicate mean ±SD. (**G**) Metagene profile plots depict the H3K27me3 ChIP signal in BoM-1833 and MDA-MB-436 cells following 72h of PHF19 knockdown (KD), centered ±25 kb around the center of H3K27me3 peaks. (**H**) Metagene profile plots depict the BoM-1833 and MDA-MB-436 cells following 72h of PHF19 knockdown (KD), clustered using k-means based on signal intensity changes in control (NT) and PHF19 knockdown (PHF19) conditions.

To determine whether PHF19 directly regulates local H3K27me3 deposition, we performed spike-in normalized H3K27me3 ChIP-seq in BoM-1833 and MDA-MB-436 cells following PHF19 depletion (Figure 5G-H). Consistent with the western blot results, metagene analysis did not reveal major shifts in H3K27me3 peak distribution upon PHF19 knockdown (Figure 5G). K-means clustering further confirmed that many H3K27me3 sites remained unaffected by PHF19 depletion. However, a subtle increase in H3K27me3 levels was observed at the highest methylated regions (strongest for cluster 1), in both cell lines (Figure 5H). Together, these findings suggest that PHF19 primarily functions as a scaffolding factor under our conditions, ensuring the spatial compartmentalization of PRC2 domains rather than modulating enzymatic activity.

### PHF19 clustering positively regulates TNBC cell motility

To investigate the functional role of PRC2-mediated PHF19 clustering in TNBC cells, we carried out global transcriptome analysis by RNA-seq in BoM-1833 and MDA-MB-436 cells following 72 hours of PHF19 depletion (Figure 6A). Differential expression analysis identified 637 downregulated and 498 upregulated genes (log_2_FC > 0.5, FDR < 0.05) in MDA-MB-436 cells. BoM-1833 cells exhibited a more modest transcriptional response, with 198 upregulated and 194 downregulated genes. Interestingly, the overlap between these differentially expressed genes and cluster 1 - H3K27me3-associated genes from ChIP-seq analysis was low (Figure 6B), supporting the hypothesis that under our experimental conditions, PHF19 influences gene expression primarily through indirect mechanisms rather than direct changes in H3K27me3 promoter methylation.

**Figure 6:**
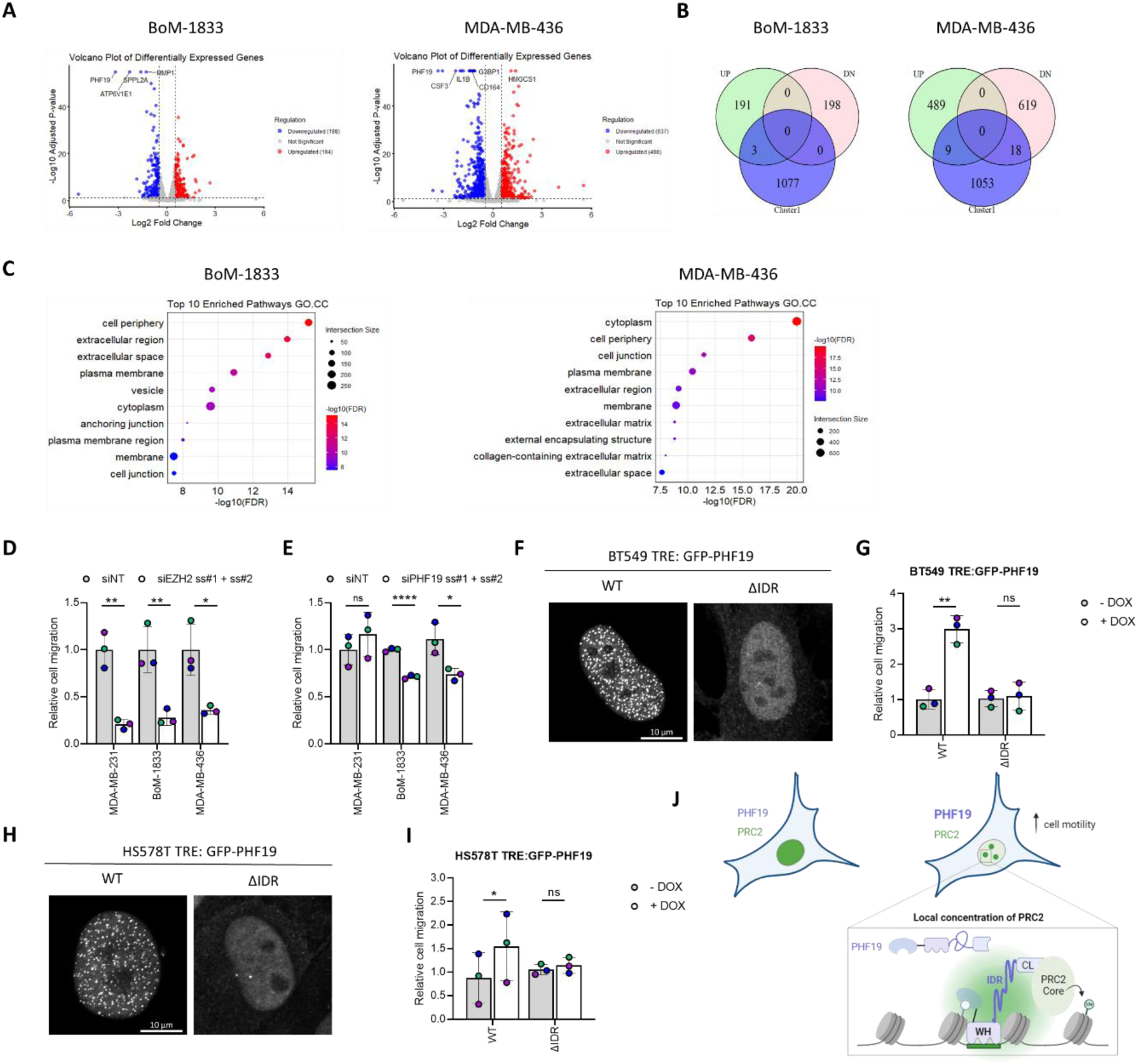
PHF19 promotes the motility of TNBC cells in an IDR dependent manner. **(A)** Volcano plots illustrate the effect of PHF19 depletion on gene expression in BoM-1833 and MDA-MB-436 cells. The RNA was isolated 72 h after PHF19 knockdown. Significantly upregulated genes (padj < 0.05, log2FC > 0.5) are highlighted in red, while significantly downregulated genes (padj < 0.05, log2FC < −0.5) are shown in blue. All other genes are represented in gray. **(B)** Venn diagram shows overlap of up-(UP) and downregulated (DN) genes from mRNA-seq analysis with genes associated with cluster 1 of the H3K27me3 ChIP-seq. **(C)** Dot plots illustrate enriched pathways from GO.CC associated with differentially regulated genes from mRNA-Seq analysis shown in (A). Color intensity represents the significance of enrichment, while dot size indicates the number of overlapping genes within each pathway. (**D-E**) MDA-MB-231, BoM-1833 and MDA-MB-436 cells were transfected with the indicated siRNAs. Transwell migration assays were performed 96h later using a serum gradient. The data are reported relative to the mean values of the siNT control. Data represent measurements from n = 3 biological replicates. Biological repeats are color coded. Statistical significance was determined via unpaired t-test. (D) p= 0.0022 (MDA-MB-231); 0.0086 (BoM-1833); 0.0156 (MDA-MB-436). (E) p= 0.3781 (MDA-MB-231); <0.0001 (BoM-1833); 0.0217 (MDA-MB-436). (**F, H**) Representative confocal microscopy image of BT549 (F) and HS578T (H) cell lines with stable, inducible expression of GFP-PHF19 and GFP-PHF19 ΔIDR, respectively. The cells were fixed 24h after doxycycline addition. Image acquisition and display settings are identical among conditions that are to be directly compared. Scale bar: 10µm. (**G, I**) Transwell motility assays were performed using BT549 (G) and HS578T (I) cells with stable, inducible expression of GFP-PHF19 and GFP-PHF19 ΔIDR, respectively. 24h after doxycycline treatment, the cells were subjected to motility assays towards a serum gradient. Data represent measurements from n = 3-4 biological replicates. Biological repeats are color coded. The data are reported relative to the mean values of the - dox control. Statistical significance was determined via paired t-test. (G) p= 0.0046 (WT); 0.5463 (ΔIDR). (I) p= 0.0271 (WT); 0.3439 (ΔIDR). (**J**) Schematic representation of the molecular pathway uncovered in this work. WH: Winged helix domain, IDR: Intrinsically disordered region, CL: Chromolike domain. The figure was created using Biorender.

Notably, GO cellular compartment enrichment analysis (Figure 6C), revealed that pathways altered upon PHF19 depletion were associated with the extracellular space, the collagen matrix and cell junctions. In both BoM-1833 and MDA-MB-436 cells, we observed a downregulation of genes encoding actin-remodeling proteins, such as *Tropomyosin 4 (TPM4)* ^54^, *ENAH* ^55^, and *Rho Guanine Nucleotide Exchange Factor 4* (*ARHGEF4*) ^56^. Additionally, expression of *ADAMTS6*, a metalloprotease that modulates focal adhesion stability and transforming growth factor β (TGFβ) bioavailability ^57^, was also reduced. Collectively, these data point towards a role of PHF19 in regulating the adhesion and migration of TNBC cells.

To test this hypothesis, we performed directed cell motility assays using a serum gradient, with the underside of the transwell coated with collagen. Consistent with previous studies ^58,59^, EZH2 knockdown significantly impaired migration in MDA-MB-231, BoM-1833, and MDA-MB-436 cells (Figure 6D), confirming the broad importance of PRC2 activity in supporting motility in these cellular systems. Strikingly, PHF19 depletion selectively dampened migration in BoM-1833 and MDA-MB-436 cells, both of which exhibit PRC2 bodies, whereas no significant effect was observed in MDA-MB-231 cells, where the PRC2 core subunits localize largely diffusely (Figure 6E). Within the timeframe of our assay, the viability and proliferation of BoM-1833 and MDA-MB-436 cells was unaffected by PHF19 depletion confirming that the observed effects were specific to motility (Figure S6). These findings suggest that disrupting PRC2 compartmentalization locally via PHF19 depletion can selectively impinge on cellular functions without the need to disrupt PRC2 enzymatic function globally.

To evaluate the direct contribution of PHF19 clustering to cell behavior, we engineered BT549 cells with doxycycline-inducible expression of either WT or ΔIDR PHF19. Western blot analysis validated that both GFP-tagged protein variants were expressed at comparable expression levels (Figure S7). Furthermore, microscopy analysis confirmed that while WT PHF19 formed visibly distinct nuclear bodies, the ΔIDR mutant displayed a fully diffuse localization throughout the nucleus (Figure 6F). This underscores the importance of the IDR for PHF19 clustering. Notably, transwell migration assays revealed that WT GFP-PHF19 expression significantly enhanced cell motility, whereas cells expressing the ΔIDR mutant exhibited no change compared to the no-dox control (Figure 6G). These observations were further validated in HS578T cells, where WT PHF19 formed clusters and promoted cell migration, while the ΔIDR mutant was diffuse and failed to do so (Figures 6H–I and Figure S7).

## Discussion

PRC2 is a master regulator of epigenetic silencing and chromatin organization, playing a pivotal role in the maintenance of cellular identity programs during normal development ^1,60,61^. Its deregulation has been implicated in numerous diseases, including cancer, where PRC2 subunit overexpression, mutation or chromatin mis-targeting contributes to aberrant transcriptional programs and disease progression ^2,62^.

Here, we uncover a previously unrecognized layer of PRC2 regulation in cancer cells. By integrating high-resolution imaging, spatial proteomics, chromatin profiling, and targeted perturbation experiments, we find that sub-stoichiometric accessory subunits actively shape the spatial organization of PRC2 domains within the nuclei of cancer cells. Specifically, we show that PHF19 modulates the compartmentalization of PRC2 into micron-sized nuclear bodies, and that this higher-order spatial arrangement has functional consequences on the cellular phenotype. While PCL proteins are known to modulate the recruitment of PRC2 to chromatin as well as its enzymatic activity ^2,11,38^, our findings are the first to show that PHF19 directly governs the spatial organization of PRC2 within the nucleus of cancer cells at the micron-scale domain level. In the context of TNBC, our work furthermore expands the paradigm of PRC2 regulation at the level of nuclear architecture, moving beyond prior studies on EZH2 overexpression or non-canonical functions in the cytoplasm ^30,62^.

The PRC2 clusters identified in our work share structural and functional similarities with PcG bodies previously described in mammalian embryonic stem cells. These act as repressive chromatin hubs and have been most extensively studied in relation to PRC1 organization ^12,15,18,21,38,48^. However, in contrast to PcG bodies, which typically disperse upon differentiation, resolving into smaller chromatin domains of less than 200 nm ^15,16,24^, we find that this higher order PRC2 spatial organization persists in TNBC cells. Given that PHF19 is downregulated during normal differentiation ^39,63^ but is upregulated in TNBC, our findings suggest that cancer cells hijack a developmental mechanism of PRC2 compartmentalization to regulate disease promoting pathways. Notably, although PcG bodies are detected across all these systems ^16,64^, whether their composition and molecular interactions are conserved across cell types and physiological contexts remains unknown. Optoproteomics could help identify commonalities and variations in PcG body composition across different cellular states. Complemented with spatial genomic approaches such as photoselective sequencing ^65^, this could provide valuable and unbiased insights into this fundamental regulatory aspect of Polycomb biology.

Biochemically, we found that PHF19 can drive PRC2 clustering *de novo*, owing to its intrinsic ability to form nuclear bodies. Our analyses revealed that this property emerges from several domains in PHF19, including its DNA-binding domain, the SUZ12 interaction interface as well as an intrinsically disordered region (IDR), which we newly mapped in this work. A role of IDRs in the subcellular compartmentalization of PCL proteins was also recently proposed by Lu and Li (2003) ^49^ who identified an IDR in PHF1 and found that this is a critical mediator of PRC2 organization via phase separation. However, while the study on PHF1 primarily relied on overexpression systems, our work leveraged physiological expression levels of PHF19, uncovering its spatial organization in cellular models of TNBC without artificial overexpression. To our knowledge, this work is the first to show that PHF19 mediates the assembly of PRC2 into microscopically distinct nuclear bodies under endogenous conditions and which links this spatial organization to functional outcomes relevant for cancer cell biology. Intriguingly, the amino acid sequence within the PHF19 IDR region is poorly conserved in PHF1 and MTF2 ^66^. This might explain why the other two PCL family members, despite being expressed in BoM-1833 and MDA-MB-436 cells, cannot compensate for the role of PHF19 in PRC2 compartmentalization. Together, these findings contribute to the emerging picture of PCL proteins exhibiting substantial functional differences ^3,67^. This conclusion is further supported by our TCGA data analysis, which showed selective upregulation of PHF19—but not PHF1 or MTF2—in TNBC.

A positive contribution of PHF19 to cancer cell behavior was previously described in multiple myeloma and prostate cancer ^68,69^. In melanoma models, PHF19 depletion hindered tumorigenesis and disrupted the H3K27me3 landscape by leading to a loss of broad H3K27me3 domains. Conversely, in prostate cancer, PHF19 depletion dampened cell motility but resulted in increased H3K27me3 levels. The latter was attributed in part to MTF2 upregulation, which functionally compensated for the loss of PHF19 ^69^. Intriguingly, the consequences of PHF19 on H3K27me3 distribution seem to depend on the cellular context. For instance, in mouse embryonic stem cells, PHF19 depletion resulted in a global reduction of H3K27me3 ^38^ while in hematopoietic stem cells, PHF19 knockout caused a global increase in H3K27me3 with a notable accumulation in blood lineage-specific genes ^39^. In our hands, PHF19 knockdown caused fragmentation of large H3K27me3 domains, as revealed by microscopy. However, we did not observe significant alterations in H3K27me3 deposition, by neither western blot or spike-in normalized ChIP-seq. Beyond the specific cellular context of TNBC, a key distinction of our study is the acute nature of PHF19 perturbation, which spans only a few days. This allows us to capture changes in the short-term cellular response before long-term compensatory mechanisms take effect. Whitin this time line, our data suggests that the regulation of gene expression by PHF19 is indirect, probably through local reorganizations in higher order chromatin structure, rather than through direct transcriptional control via CpG promoter methylation. Notably, also ^69^ reported minimal overlap between gene expression changes and direct PHF19 binding. This suggest that the role of PHF19 in cell behavior, through larger scale chromatin re-organization, could be partly conserved in other cellular systems beyond the ones explored in our work.

Beyond PHF19 and PCLs, which define the PRC2.1 complex, PRC2 can also associate with AEBP2 and JARID2 to build the PRC2.2 subtype ^4^. The establishment and maintenance of H3K27me3 methylation patterns involves the cooperative behavior between PRC2.1 and PRC2.2, which act through both independent and synergistic mechanisms. Indeed, studies in mouse embryonic stem cells have shown that while the loss of either PRC2.1 or PRC2.2 alone was insufficient to deplete H3K27me3 globally, the combined depletion of both complexes disrupted core PRC2 targeting and stable tethering to Polycomb target genes, resulting in genome-wide PRC2 mislocalization ^70^. For the future, it would be interesting to explore the crosstalk between PRC2.1 and PRC2.2 accessory subunits in PRC2 spatial organization and function in TNBC cells.

Collectively, our data support a model in which PHF19 governs the subnuclear organization patterns of PRC2 in TNBC cells (Figure 6J). We propose that PHF19 integrates recognition of DNA sequence motifs with specific chromatin features, such as PRC2 occupancy and H3K36 methylation, to target specific genomic regions. Once bound, PHF19 promotes PRC2 clustering at these sites via its IDR, facilitating multivalent interactions that amplify local PRC2 enrichment. These nuclear hubs drive the formation of large chromatin domains, which, through their spatial organization, regulate gene expression programs central to cellular behavior such as cell motility. We hypothesize that by concentrating PRC2 activity within defined nuclear hubs, this organization may improve the robustness and fidelity of Polycomb-controlled gene expression programs, a property hijacked in cancer to foster aggressive cellular phenotypes.

By uncovering the unique role of PHF19 in PRC2 spatial organization, our study advances the understanding of epigenetic regulation in both normal development and disease. In cancer, these findings open new avenues for therapeutic intervention, including strategies targeting PRC2 clustering to disrupt cancer-specific gene expression programs. Furthermore, PHF19 expression could serve as a biomarker for patient stratification in TNBC, providing a foundation for precision medicine approaches. Together, our work underscores the importance of spatial biology in cancer epigenetics and highlights the therapeutic potential of targeting context-dependent PRC2 regulation.

### Limitations of the study

While our study provides significant mechanistic insights into PRC2 regulation in the cancer context, several limitations warrant consideration. First, the reliance on a limited number of TNBC cell lines may not fully capture the heterogeneity seen in patient tumors. Additionally, while we demonstrated the role of PHF19 in PRC2 clustering and cell motility *in vitro*, its relevance to tumor progression and metastasis *in vivo* remains unexplored and should be investigated in the future.

## Materials and methods

### Cell culture

MDA-MB-231, BT549 (obtained from CLS Cell Lines Service GmbH, Eppelheim, Germany), BoM-1833 (kindly provided by Joan Massagué, Memorial Sloan Kettering Cancer Center, USA), MDA-MB-436 (kindly provided by the Institute of Clinical Pharmacology, Stuttgart, Germany), HS578T (kindly provided by Bernhard Lüscher, RWTH Aachen, Germany) and LentiX (kindly provided by Philipp Rathert, Institute of Biochemistry and Technical Biochemistry, University of Stuttgart, Germany) cells were cultured in DMEM (Invitrogen) supplemented with 10% FCS and grown under sterile conditions in a humidified atmosphere of 5% CO_2_ at 37°C. All cell lines were authenticated, tested negative for Mycoplasma (Lonza, LT07-318) and were kept in culture for no longer than 2 months.

### Antibodies

The following antibodies were used in this study: rabbit anti-EZH2 (Cell Signaling Technology Cat# 5246, RRID:AB_10694683, 1:200 in IF and 1:1000 in WB), rabbit anti-GAPDH (Sigma-Aldrich Cat# G9545, RRID:AB_796208, 1:5000 in WB), rabbit anti-GFP (Cell Signaling Technology Cat# 2956, RRID:AB_1196615, 1:1000 in WB), rabbit anti-H3 (Cell Signaling Technology Cat# 4620, RRID:AB_1904005, 1:5000 in WB), mouse anti-H3K27me3 (Diagenode Cat# C15200181, RRID:AB_2819192, 1:250 in IF, 1:500 in WB), rabbit anti-H3K27me3 (Active Motif Cat# 39155, RRID:AB_2561020, 5 µg per ChIP-seq reaction), rabbit anti-MTF2 (Proteintech Cat# 16208-1-AP, RRID:AB_2147370, 1:1000 in WB), rabbit anti-PHF1 (Abcam Cat# ab184951, RRID:AB_2861270, 1:1000 in WB), rabbit anti-PHF19 (Cell Signaling Technology Cat# 87162, 1:250 in IF and 1:1000 in WB) and rabbit anti-SUZ12 (Cell Signaling Technology Cat# 3737, RRID:AB_2196850, 1:200 in IF and 1:1000 in WB).

HRP-labeled secondary goat anti-mouse and anti-rabbit IgG antibodies were purchased from Dianova (Hamburg, Germany), Alexa-Fluor-labeled secondary IgG antibodies were from Invitrogen. DAPI was from Sigma-Aldrich (5 µg/mL in IF).

### DNA constructs

The GFP-PHF19 plasmid was generated using Gibson assembly (NEB). The PHF19 insert was amplified by PCR from pCR8-PHF19 (a gift from Adrian Bracken, Addgene #68864) using the primers Fw: 5’-GAGAATCGAGCTCTGGATCCAG-3’ and Rev: 5’-GTCAGTAAGGGGTGGTCCCTTCC-3’ and subcloned into pEGFP-C1 (Clontech), which had been linearized by PCR using Fw: 5’-GGACCACCCCTTACTGACCCGGGATCCACCGGATCTAG-3’ and Rev: 5’-GATCCAGAGCTCGATTCTCTGCAGAATTCGAAGCTTGAGCTCG-3’. The GFP-PHF19ΔIDR truncation construct was generated through QuickChange mutagenesis of the GFP-PHF19 plasmid using the primers Fw: 5’-TTTGTCAGGCAGCAGCTTCC-3’ and Rev: 5’-GATGACTCATCCCTGTCCCAC-3’. Similarly, the GFP-PHF19 K331/332A mutant was generated using QuickChange mutagenesis of the GFP-PHF19 plasmid with the primers Fw: 5’-GGCAAGGAGATCGCGGCGAAGAAGTGCATCTTCCGCC-3’ and Rev: 5’-GATGCACTTCTTCGCCGCGATCTCCTTGCCGCAGAG-3’.

The mScarlet-IDR plasmid was generated using Gibson assembly (NEB), whereby the IDR domain of PHF19 (356aa–528aa) was amplified by PCR from GFP-PHF19 using the primers Fw: 5’-GATCTCGAGCTCAAGCTGGACTGCTGCCAAATGAGAACAG-3’ and Rev: 5’-CGACTGCAGAATTCGATTATTCACTGATGCTCTCAAATGTGTGG-3’. The fragment was then subcloned into mScarlet-C1 (Clontech), which was linearized by PCR using Fw: 5’-AGCTTGAGCTCGAGATCTGAGTC-3’ and Rev: 5’-TAATCGAATTCTGCAGTCGACGG-3’.

The mScarlet-IDR-Chromolike plasmid was also generated by Gibson assembly (NEB). The IDR-Chromolike region of PHF19 (356aa – 580aa) was amplified by PCR from GFP-PHF19 using the primers Fw: 5’-GATCTCGAGCTCAAGCTGGACTGCTGCCAAATGAGAACAG-3’ and Rev: 5’-CGACTGCAGAATTCGATTAGTCAGTAAGGGGTGGTCCCTTCC-3’. The amplified fragment was subcloned into mScarlet-C1 (Clontech), which was linearized by PCR using Fw: 5’-AGCTTGAGCTCGAGATCTGAGTC-3’ and Rev: 5’-TAATCGAATTCTGCAGTCGACGG-3’.

The doxycycline-inducible GFP-PHF19 and GFP-PHF19ΔIDR lentiviral transduction plasmids were generated using Gibson assembly (NEB), whereby the inserts were amplified by PCR from GFP-PHF19 and GFP-PHF19ΔIDR, respectively, using the primers Fw: 5’-GATCGCCTGGAGAATTGGCTAGATGGTGAGCAAGGGCGAG-3’ and Rev: 5’-GTACAACGCGTCTGCAGCCTAGTCAGTAAGGGGTGGTCCCTTC-3’. The fragments were then subcloned into pCW57-MCS1-P2A-MCS2 (Neo) (Addgene #89180), which was linearized with AvrII and NheI (NEB).

The identity of all constructs was validated by full plasmid sequencing (Microsynth Seqlab, Göttingen, Germany).

### Immunofluorescence staining and confocal microscopy

Cells were grown on glass coverslips pre-coated with 10 μg/ml collagen R (Serva) and fixed 24 – 48 h later for 10 min at room temperature using 4% (w/v) paraformaldehyde. After washing with PBS supplemented with Ca^2+^ and Mg^2+^, the cells were incubated for 15 min with 150 mM glycine in PBS followed by permeabilization for 5 min with 0.1% (v/v) Triton X-100 in PBS. Blocking was performed with 5% (v/v) goat serum (Invitrogen) in PBS containing 0.1% (v/v) Tween-20. Fixed cells were incubated overnight at 4°C with primary antibodies diluted in blocking buffer. Following three washing steps with PBS, the cells were incubated for 1 h at room temperature with Alexa-Fluor®-(488, 555 or 647)-labelled secondary antibodies diluted in blocking buffer. Nuclei were counterstained with DAPI, the actin cytoskeleton was labeled using Phalloidin-633 (ThermoFisher Scientific) and the coverslips were mounted in Molecular Probes™ ProLong™ Gold Antifade mountant (ThermoFisher Scientific). Imaging was performed on an LSM980 Airyscan 2 (Carl Zeiss, Oberkochen, Germany) equipped with a Plan-Apochromat 63x/1.40 DIC (Carl Zeiss) oil immersion objective using 405-, 488-, 561-nm laser excitation. For each set of replicates, images were acquired with the same laser and confocal settings. Maximum intensity projections, linear adjustments of brightness and contrast, and analysis of mean fluorescence intensity (MFI) of regions of interest (ROI) was performed with the ZEN software (Zeiss) or Fiji.

### Image analysis

To measure the diameter of PRC2 and PHF19 bodies (Figure 1C & I, Figure 3J), raw imaging files were processed using Fiji. The diameters were determined manually using the line function.

To determine the percentage of cell nuclei containing PRC2 core subunits, PHF19, and mScarlet-IDR bodies (Figure 1D & J, Figure 2C, Figure 3D & O, Figure 4G), raw image files were analyzed in Fiji. Individual cells were manually inspected, and those exhibiting more than one distinct cluster of the protein of interest—clearly distinguishable from the diffuse nucleoplasmic signal and at least 1 µm in size—were classified as positive. No size cut-off was applied for mScarlet-IDR bodies.

To quantify PRC2 clusters in cells with transient PHF19 overexpression (Figure 3P), raw image files were processed in Fiji. Images were converted to 8-bit, and color channels were separated. A Gaussian blur filter with a sigma (radius) of 3.00 was applied, followed by manual thresholding to distinguish visible PRC2 clusters from background granularity. Using the Analyze > Analyze Particles function, particles with a minimum size of 0.25 µm and a circularity of at least 0.5 were counted. Nuclei containing more than three PRC2 clusters were classified as PRC2 cluster-positive. The percentage of PRC2 cluster-positive nuclei in transfected cells was compared to non-transfected cells and plotted accordingly in Prism (GraphPad).

To determine the relative enrichment of GFP-PHF19 and its variants at H3K27me3 foci (Figure 4E & I), raw image files were analyzed manually in Fiji. H3K27me3 staining was used to define the cluster ROI, which was then used to generate a mask for measuring the mean MFI of the GFP channel. Background fluorescence in the nucleoplasm was determined by averaging the GFP intensity from three arbitrarily chosen ROIs placed adjacent to the H3K27me3 cluster of interest.

To assess whether PHF19 knockdown affects the appearance of H3K27me3 clusters in MDA-MB-436 cells (Figure 5F), the number of clusters per nucleus was quantified in Fiji using an automated pipeline. First, a mask was created to identify cell nuclei based on DAPI staining, excluding the background from the analysis. To reduce noise from staining granularity, a Gaussian blur filter with a sigma (radius) of 4 was applied. A threshold was then set to distinguish H3K27me3 clusters from the nuclear background. Clusters larger than 0.3 µm were counted using the “Analyze Particles” function, and cells were categorized into four groups based on the number of clusters per nucleus: 0 clusters, 1-3 clusters, 4-6 clusters, and more than 6 clusters. Due to the high granularity of the H3K27me3 signal, this pipeline could not be applied to analyze images obtained with BoM-1833 cells. Instead, for this cell line, the percentage of nuclei positive for H3K27me3 accumulation was quantified manually in Fiji (Figure 4D). Nuclei were classified as positive if they exhibited visible H3K27me3 foci larger than 0.5 µm.

To assess the relationship between GFP-PHF19 expression levels and the number of GFP-PHF19 clusters per nucleus (Figure 4B), raw image files were analyzed in Fiji. A mask was created on 8-bit images to identify cell nuclei based on DAPI staining. The mean fluorescence intensity (MFI) of GFP was extracted within the nuclear region of interest (ROI). Additionally, the Analyze Particles function was used to count GFP clusters larger than 0.3 µm.

Line profiles (Figure 4D & F) were generated using Fiji. To this end, a line was drawn through the region of interest, and its position was saved. This line was then transferred to all channels of interest. The MFI along the line was extracted using the “Analyze > Plot Profile” function and plotted for analysis in Prism (GraphPad).

### Cell lysis and immunoblotting

Cells were lysed in lysis buffer [150 mM Tris (pH 7.5), 500 mM NaCl, 10% glycerol, 10 mM MgCl_2_, 1 mM EGTA, 1% Triton-X-100, 0.1% SDS, 0.5% sodium deoxycholate, 0.5 mM PMSF, 1 mM sodium orthovanadate, 10 mM sodium fluoride, and 20 mM β-glycerophosphate plus Complete protease inhibitors without EDTA (Roche)] by sonication with an EpiShear Probe Sonicator (Active Motif). Each sample was sonicated with 2 pulses of 30 s at an amplitude of 40%, a pause of 30 s was used after each pulse. Subsequently, the samples were incubated at 4°C for 15 min while rolling end over end, followed by centrifugation for 10 min at 16,000 g and 4°C. The protein concentration of the clarified lysate was determined by Bio-Rad DC protein assay. Lysates were loaded on 4–12% NuPAGE® Novex Bis–Tris gels (Invitrogen) and transferred to nitrocellulose membranes (iBlot®Gel Transfer Stacks; Invitrogen). Membranes were blocked with 0.5% blocking reagent (Roche) in PBS containing 0.05% Tween-20 and incubated with primary antibodies, followed by HRP-labeled secondary antibodies for ECL-based (Pierce, Rockford, IL) visualization. The chemiluminescence signal was detected using an AmershamTM Imager 600 device (GE Healthcare) followed by quantification of the 16-bit images in the linear range using the inbuilt ImageQuant TL 8.1 software.

### RNA interference and plasmid transfection

For transient knockdowns, the cells were reverse transfected using Lipofectamine™ RNAiMAX (Invitrogen) at a final concentration of 5nM total siRNA, according to manufacturer’s instructions. In experiments involving siRNA mixtures, the individual siRNAs were mixed 1:1. The cells were used for experiments at 72-120 h post transfection as described in the figure legends. The siRNAs used were: negative control siRNA (siNT, ON-TARGETplus® non-targeting control pool D-001810-10 from Dharmacon), siPHF19 (Silencer® Select PHF19 s25161 (ss#1) and s25160 (ss#2) from ThermoFisher Scientific) and siEZH2 (Silencer® Select EZH2 s4917 (ss#1) and s4918 (ss#2) from ThermoFisher Scientific). For plasmid transfections, LipofectamineLTX (ThermoFisher Scientific) was used according to manufacturer’s instructions.

### Stable cell line generation

Lentiviral particles were produced using LentiX HEK293T cells (a gift from Philipp Rathert, University of Stuttgart), which were transfected with the packaging plasmid psPAX2 (a gift from Didier Trono, Addgene #12260) and pCMV_VSV-G (a gift from Bob Weinberg, Addgene #8454) as well as the pCW57 plasmid containing the transgene of interest. The supernatant was collected over 24 h, passed through 0.4 µm filters and added to the target cells. Transduced cells were selected with 500 µg/mL G418 for at least 14 days before being used for experiments. Transgene expression was induced by addition of 2 µg/mL Doxycycline (Sigma).

### Boyden chamber assays

The cells were treated with siRNAs or doxycycline as indicated in the figure legends, harvested and re-suspended in media containing 0.5% FCS. The underside of the transwells (8.0 µm pore Size, 3422, Costar) was coated with 25 ug/ml Collagen-R (Serva) ON at 4°C. 50.000 cells were seeded in the upper transwell compartment. The media in the lower compartment was supplemented with 10% FCS. At 2.5 h (HS578T), 4 h (MDA-MB-231 and BoM-1833), 4.5 h (HS578T) and 24 h (MDA-MB-436) post seeding, the transwells were fixed with ROTI®Histofix (P087.3, Roth) for 15 min and the cells were stained with DAPI. From each transwell at least 5 different field of views were imaged using an EVOS M5000 and the average number of cells per image calculated in Fiji. To this end, the images were converted to 8-bit format. Background noise was removed, and an automatic threshold was applied to define the nuclei. This threshold was used to create a mask, and adjacent nuclei were separated using the “Watershed” function. The “Analyze Particles” plugin was used to count nuclei that were larger than 8 µm and had a circularity factor of at least 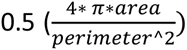. Nuclei that were cut off at the edges of the images were excluded from the analysis.

### Cell viability measurements

The cells were treated with siRNAs as indicated in the figure legends, and seeded in triplicates in 12-well plates, one triplicate set was measured per time point. To determine viability, the cells were harvested at the indicated intervals and were stained with trypan blue (Invitrogen). Both viable and non-viable cell counts were measured using the Countess 3 automated cell counter (Invitrogen). The viability percentage was calculated based on the proportion of unstained (live) cells relative to the total cell count.

### Cell culture for Microscoop® photolabeling

BoM-1833 cells were cultured in RPMI 1640 medium (11875093, Gibco, Thermo Fisher Scientific, USA) containing 10% FBS (A52567-01, Gibco, Thermo Fisher Scientific, USA), antibiotics (15140-122, Gibco, Thermo Fisher Scientific, USA), and maintained in a 37℃ humidified incubator with 5% CO_2_. To prepare the samples, approximately 4.5 × 10⁵ cells were seeded at 80% confluency in a one-well chamber slide (C1-1.5H-N, Cellvis, USA). For each biological replicate, five chamber slides were used for photolabeling (PL), while another five chamber slides served as unlabeled controls (UL).

### Immunofluorescence staining followed by Microscoop® photolabeling

BoM-1833 cells were fixed by incubating with 2.4% paraformaldehyde solution for 10 mins RT, rinsed 2 times with PBS and then permeabilized with 0.5% PBSTx (PBS containing 0.5% Triton X-100) for 10 mins at RT. After 3 PBS washes, the cells were blocked with 3% BSA diluted in PBS-Tx for 1 hour at RT. Samples for Microscoop® photolabeling were prepared by Syncell using the Synlight-RichTM kit (SYN-PU0106, Syncell, Taiwan) according to the manufacturer’s instruction. Briefly, the cells were blocked with Block 1 and Block 2 blocking reagents for 30 and 15 min respectively. The cells were then incubated with the rabbit anti-EZH2 antibody (5246, Cell signaling, USA) for 4 hours at RT, washed 3 times with PBST for 5 min and then incubated with Alexa Fluor™ 647 secondary antibody (A-21245, ThermoFisher, USA) for 2 hours. The image processing function in Microscoop® was used to generate a mask for EZH2 clusters, which automatically guided the photolabeling process within the masked regions across the entire cell chamber slide. Upon completion of the photolabeling process, the cells were quenched with the quencher in Synlight-RichTM kit (SYN-RI0106, Syncell, Taiwan) for 3 times wash, each for 5 minutes. Subsequently, the cells were thoroughly rinsed with PBS-Tx to minimize interference from unreacted probes during the pulldown procedure. Unlabeled samples were processed in parallel and were used as control.

To verify the biotinylation efficiency, the cells were stained with the “Verify” reagent in the Synlight-RichTM kit (SYN-RI0106, Syncell, Taiwan) for 1 hour at room temperature. The images were obtained by epi-fluorescence microscopy (TE2000, NIKON, Japan) or confocal (LSM880, ZEISS, Germany). For confocal imaging, the DNA was counterstained with Hoechst-33342 (H3570, Thermo Fisher Scientific, USA).

### Sample immunoprecipitation for mass spectrometry

Sample processing for mass spectrometry was prepared by Syncell using the SynpullTM reagent kit (SYN-PU0106, Syncell, Taiwan) according to the manufacturer’s instruction. Briefly, the photolabeled and unlabeled cells were scraped with scraping buffer and collected in separate tubes. After centrifugation, most of the supernatant was discarded while cell pellet and approximately 100 μl of buffer was retained. 60 μl lysis buffer was added to the pellet and the cells were lysed by sonication for 60 cycles (1s on / 2s off) at 30% amplitude with a probe-based sonicator (Q-125, Qsonica, USA). The sample was spun down and then heated at 99℃ for 1 hour. Solution D was added to the cooled down samples, followed by thorough mixing and incubation at room temperature for 30 minutes in the dark. After incubation, the samples were centrifuged and the supernatant transferred into new tubes to determine the protein concentration using the Pierce 660 nm protein assay (IDCR method). Equal amount (200 µg) of proteins was used as IP input for both the 2P (labeled) and CTL (unlabeled) group. The proteins were diluted five-fold with solution E and streptavidin magnetic beads were added to the lysate, followed by incubation with rotation for 1 hour at room temperature to pull down biotinylated proteins. Protein-bound beads were washed four times using the buffers provided with the kit. The beads were finally resuspended in wash buffer and subsequently sonicated for 10 seconds using a ultrasonic water bath. The supernatant was then discarded and the beads were resuspended in 20 µL digestion buffer to conduct on-beads digestion. After 2 h incubation at 37℃ with 35° shaking using a Intelli-MixerTM (RM-2S, ELMI, Latvia), the supernatant was transferred into the proteomics tubes provided in the kit and the incubation was continued at 37℃ for 16 h with 35° shaking. Solution L was added to each sample to stop reaction and a desalting step was subsequently performed. The desalted peptides were concentrated by a SpeedVac concentrator until completely dried.

### Detection of the immunoprecipitated product by data-dependent acquisition mass spectrometry

The pull-down biotinylated proteins were detected using data-dependent acquisition (DDA) mass spectrometry. Liquid chromatography-tandem mass spectrometry (LC-MS/MS) was performed by Syncell using an UltiMate 3000 RSLCnano system (Thermo Fisher Scientific, USA) coupled to an Orbitrap Fusion Lumos mass spectrometer (Thermo Fisher Scientific, USA). The desalted peptides were resuspended in 0.1% formic acid in water and loaded onto a PepMap™ 100 C18 HPLC column (2 µm, 100 angstrom, 75 µm × 25 cm; Thermo Fisher Scientific, USA), and peptides were eluted over 120 min gradients. The full MS spectra ranging from m/z 375–1500 were acquired at a resolving power of 120,000 in Orbitrap, an AGC target value of 4×10^5^, and a maximum injection time of 50 ms. Fragment ion spectra were recorded in the top-speed mode at a resolving power of 30,000 in Orbitrap using a data-dependent method. Monoisotopic precursor ions were selected by the quadrupole using an isolation window of 1.2, 0.7, 0.4 Th for the ion with 2+, 3+, 4–7 charge states, respectively. An AGC target of 5×10^4^, maximum injection time of 54 ms, higher-energy collisional dissociation (HCD) fragmentation with 30% collision energy, and a maximum cycle time of 3 s were applied. Dynamic exclusion was set to 60s with an exclusion window of 10 ppm. Precursor ions with the charge state of unassigned, 1+, or superior to 8+ were excluded from fragmentation selection.

### Protein identification and label-free quantification

Raw data from the same batch of two-photon illumination were processed together with Proteome Discoverer (version 2.4.1.15; Thermo Fisher Scientific, USA) by Sequest HT algorithm against the UniProtKB/Swiss-Prot human protein database (version 2020.03, 20,365 entries) for feature extraction, peptide identification, and protein inference. Database search was performed as follows: tryptic peptides with up to three missed cleavages; mass tolerances of 10 ppm for peptide ions, and 0.05 Da for fragment ions; static carbamidomethylation (+57.0215 Da) on Cys residues; dynamic deamidation (+0.9840 Da) on Asp and Gln residues, oxidation (15.9949 Da) on Met residues, and acetylation on protein N-termini (+42.0106 Da). The minimal peptide length was set as 6 residues. The false discovery rate (FDR) of peptide and protein were both set as 1%. For label-free quantification, the time window for chromatographic peak alignment was set as 20 min. Peptide level data was then normalized to the total peptide intensity, and the quantification value for a given protein was derived from the sum of normalized intensities of the top three intense unique peptides belonging to that protein.

### RNA-seq library preparation and sequencing analysis

Total RNA was isolated using the NucleoSpin RNA Midi Kit (Macherey-Nagel, 740962.20) using the manufacturer’s instructions. RNA integrity was assessed by gel electrophoresis, and libraries were prepared using 1 µg of RNA with the Illumina Stranded mRNA Prep Ligation Kit (Illumina, 1000000124518 v03), following the manufacturer’s protocol. After library preparation, library concentration was measured using a Qubit Fluorometer (Invitrogen) with the Qubit 1X dsDNA High Sensitivity Kit (ThermoFisher Scientific, Q33231). Library size was assessed using TapeStation (Agilent) with the High Sensitivity D1000 Sample Buffer (Agilent, 5190-6504) and High Sensitivity D1000 ScreenTape (Agilent, 5067-5584). Paired-end 61 bp sequencing was performed at the Robert Bosch Center for Tumor Diseases using the Illumina NextSeq 2000 system.

### ChIP and ChIP-seq library preparation

Chromatin immunoprecipitation (ChIP) was performed following the protocol described in Ekstrom et al., 2025 ^71^, with some modifications. Briefly, after 72 hours of siRNA treatment, 20 million cells were collected and mixed with NIH 3T3 cells (5%) as a spike-in control per cell number. Cells were crosslinked with 1% formaldehyde for 20 minutes, followed by quenching with 1.25 M glycine. Chromatin was sheared using a Bioruptor Pico (Diagenode) for 8 cycles (30 seconds on/off) to achieve fragment sizes of approximately 200-500 base pairs. After preclearing with a 50% slurry of Sepharose beads (Cytiva, 17012001), chromatin was incubated overnight with anti-H3K27me3 (Active Motif, 39155, 5 µL/IP) antibody. Protein A Sepharose beads (Cytiva, 17078001, rabbit antibodies) were subsequently added for immunoprecipitation. The immunoprecipitated chromatin was washed sequentially with high-salt buffers and TE buffer. After washing, chromatin was reverse crosslinked, and DNA was extracted. ChIP DNA quantification was performed using Qubit.

ChIP libraries were prepared using the MicroPlex Library Preparation Kit v3 (Diagenode, C05010001), following the manufacturer’s protocol. Library fragment sizes were assessed using TapeStation with High Sensitivity D1000 Sample Buffer and High Sensitivity D1000 ScreenTape. Paired-end 61 bp sequencing was performed at the Robert Bosch Center for Tumor Diseases using the Illumina NextSeq 2000 system.

### ChIP-seq bioinformatics analysis

Paired-end sequencing reads were preprocessed using nf-core/chipseq (v2.1.0) pipeline ^72^. Bowtie2 (v2.5.2) ^73^ was used to map reads to the reference genome assembly hg38. Bigwig files were scaled ignoring blacklist regions using bamCoverage (deeptools v3.5.6) ^74^ with reads per genomic content (RPGC) normalization and scaling factors calculated as following:

1. ratio_input = spike_input / (main_input+spike_input)
2. ratio_IP = spike_IP / (main_IP+spike_IP)
3. scaling_factor_sample = ratio_input / ratio_IP
4. Normalize scaling factors by dividing all scaling factors by highest scaling factor

Scaled and normalized bigwigs for replicates were merged using bigwigAverage (deeptools v3.5.6) ^74^. MACS3 (v3.0.1) ^75^ callpeak was used to call the significant peaks with –broad –cutoff 0.05 for H3K27me3. Consensus peakset across all samples per cell line were generated by mergeBed (bedtools v2.30.0) ^76^. Given that H3K27me3 present large domains, nearby peaks were stitched within the 5kb distance. ChIP occupancy on the consensus peakset was measured by computeMatrix (deeptools v3.5.6) around the center of peaks with 25kb flanking regions. The meta profiles were generated based on the computeMatrix values using plotProfile (deeptools v3.5.6). ChIP annotation were performed using ChIPseeker package (v1.34.1) ^77^

### RNA-seq bioinformatics analysis

Paired-end sequencing reads were preprocessed using nf-core/rnaseq (v3.18.0) pipeline ^72^. Reads were first mapped to the reference genome assembly hg38 using STAR (v2.7.10a) ^78^ and then quantified using RSEM (v1.3.1) ^79^. Raw counts were used for differential gene expression analysis using R (v4.4.2) and DESeq2 (v1.46.0) ^80^. Differentially expressed genes are considered when |log2(fold change)| > 0.5, adjusted p-value < 0.05. Gene set enrichment analysis was performed using gprofiler2 (v.0.2.3) for differentially expressed genes. Volcano plots and dot plots were generated using ggplot2 package (v3.5.1).

### TCGA gene expression analysis

GEPIA2 ^81^ was used to analyze *PHF1, MTF2* and *PHF19* expression in the TCGA BRCA cohort. To this end, the “expression DYI” plugin was used and the BRCA dataset was sorted using a Log2FC cutoff of 1 and a p value cutoff of 0.01.

### Statistics

Data were analyzed and plotted using GraphPad Prism 8. Information about the statistical tests used as well as about sample size N, number of independent repeats and p values can be found in the corresponding figure legends and Table S2.

## Data availability

All data generated or analyzed during this study are included in this published article and its Supplementary Information. Source data are provided with this paper in Table S2. The EZH2 proteomics results are reported in Table S1. Uncropped Western blots are provided in Figures S8-S10. All ChIP sequencing and RNA sequencing data has been deposited in the NCBI GEO database under accession numbers: GSE291242 and GSE291243, respectively. Plasmids, cell lines and BED files generated during the study are available upon reasonable request from the corresponding author.

## Author contributions

Conceptualization: CL

Methodology: WG, CL

Investigation: NP, TL, CL, WG, NS, PT, ZN

Visualization: CL, WG, ZN

Funding acquisition: CL, ZN, MAO

Project administration: CL

Supervision: CL, ZN

Writing – original draft: CL with contributions from ZN, WG, TL and MAO

Writing – review & editing: CL, ZN, MAO.

## Acknowledgements

The RNA-Seq datasets used in GEPIA2 are based on the UCSC Xena project (http://xena.ucsc.edu), which are computed by a standard pipeline. The authors gratefully acknowledge the Technology Platform “Cellular Analytics” of the Stuttgart Research Center Systems Biology for their support and assistance in this work. We thank Raluca Tamas for critical reading of the manuscript. This work was supported by the Baden-Wuerttemberg Ministry of Science, Research and Arts by a grant to CL and by the German Federal Institute for Risk Assessment Grant Agreement Number 60-0102-01.P637 to MAO. This work was further supported by the Robert Bosch Stiftung for ZN, WG and TL.

