## Supplemental figures for "PHF19 drives PRC2 sub-nuclear compartmentalization to promote motility in TNBC cells"

- **Supplement figures and tables:**

**Figure S1:** Setup and validation of the endogenous EZH2 body photo-biotinylation assay in bone entrained TNBC cells.

**Figure S2:** Validation of the PHF19 antibody specificity for immunofluorescence in TNBC cells.

**Figure S3:** PCL family gene expression analysis across a TCGA BRCA cohort sorted by breast cancer subtype.

**Figure S4:** Quantification of PRC2 subunit protein expression in MDA-MB-436 cells with PHF19 depletion.

**Figure S5:** EZH2 but not PHF19 depletion reduces global H3K27me3 levels in TNBC cells.

**Figure S6:** PHF19 depletion does not affect cell growth and viability in TNBC cells.

**Figure S7:** Characterization of the BT549 and HS578T cell models with inducible expression of the GFP-PHF19 variants.

**Figure S8:** Uncropped Western Blot images for Figures 2 and 3.

**Figure S9:** Uncropped Western Blot images for Figure 5A.

**Figure S10:** Uncropped Western Blot images for Figures S5 and S7.

**Table S1:** Local EZH2 interactome in BoM-1833 cells. List of associated proteins enriched in all four biological replicates of the optoproteomics analysis.

**Table S2**: Primary source data and statistical testing reporting

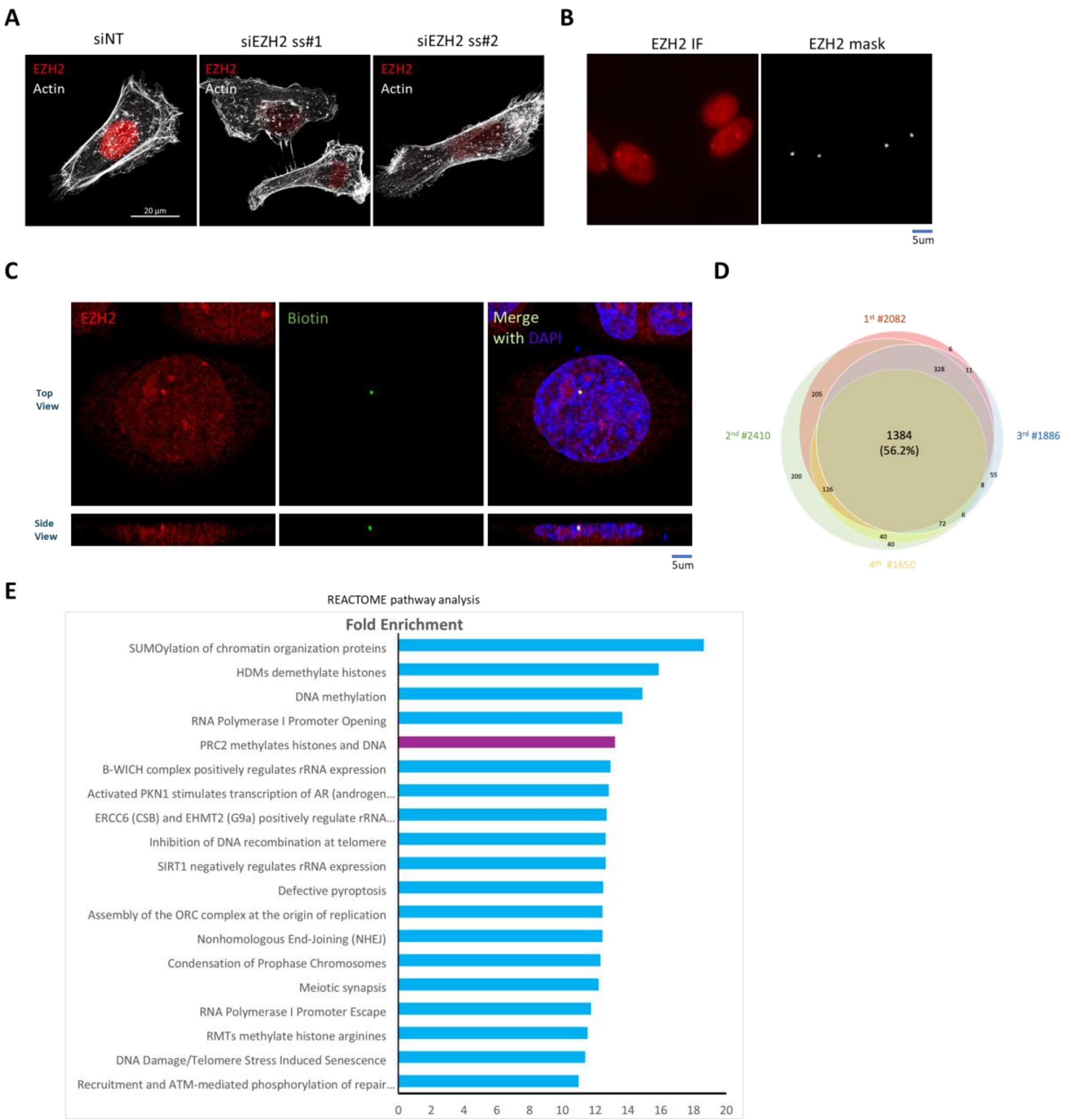


**Figure S1: Setup and validation of the endogenous EZH2 body photo-biotinylation assay in bone entrained TNBC cells.** (**A**) Validation of EZH2 antibody specificity for immunofluorescence applications. BoM-1833 cells were transfected with a non-targeting control siRNA (siNT) or two independent siRNAs targeting *EZH2*. Immunofluorescence staining was performed 96 hours post-transfection to visualize endogenous EZH2 (red) and the actin cytoskeleton (phalloidin, white). Identical image acquisition and display settings were used across conditions. Scale bar: 20 µm. (**B**) Representative example of the image segmentation process utilized for spatial EZH2 photo-biotinylation analysis. Left: Confocal microscopy image showing endogenous EZH2 immunofluorescence staining (red) in BoM-1833 cells. Right: Corresponding segmented image defining regions of interest for targeted protein labeling. Scale bar: 5 µm. (**C**) Representative confocal microscopy image of BoM-1833 cells stained for endogenous EZH2 (red) and subjected to localized biotinylation using the Microscoop® platform. Dy488-NeutrAvidin (green) was employed to detect the photo-biotinylated EZH2 protein. Nuclei are counterstained with DAPI. The figure includes a single focal plane and a side view of the 3D reconstituted image, demonstrating the spatial specificity of the biotinylation reaction. Scale bar: 5 µm. (**D**) Venn diagram summarizing the overlap in the local EZH2 proteome identified in BoM-1833 cells across four biological replicates. (**E**) REACTOME pathway analysis of targets identified from the proteomic analysis of PRC2 bodies in BoM-1833 cells, ranked by fold enrichment. The analysis was performed using DAVID on the 219 nuclear proteins enriched in the labeled versus control condition (ratio ≥ 2, -Log p-value ≥ 1.3), detected across four biological repeats with >3 unique peptides and a sequest score >0.05%. The pathway "PRC2 methylates histones and DNA" is highlighted in purple.

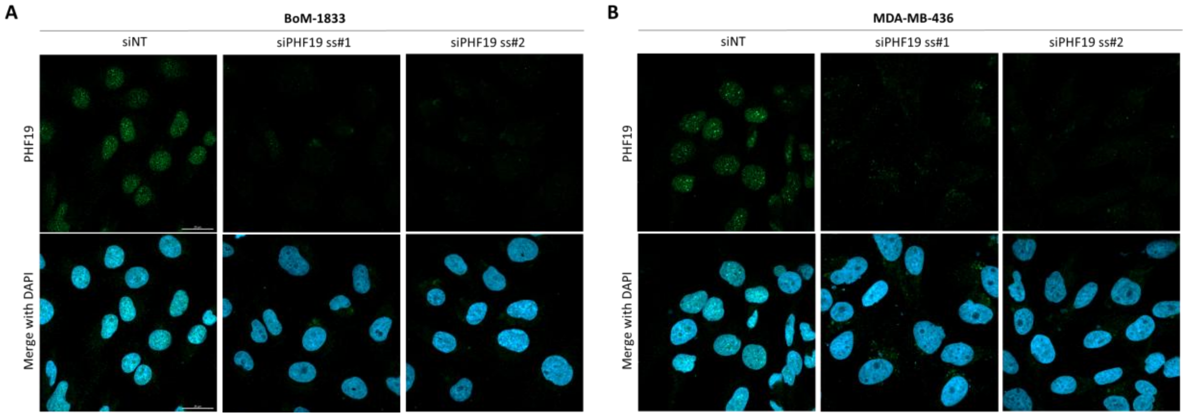


**Figure S2: Validation of the PHF19 antibody specificity for immunofluorescence in TNBC cells.** Representative confocal microscopy images of BoM-1833 (**A**) and MDA-MB-436 (**B**) cells treated with siNT or two independent siRNAs targeting PHF19. The cells were immunostained for PHF19 (green) 4 days after siRNA transfection. Nuclei were counterstained with DAPI. Identical image acquisition and display settings were used across conditions that are to be directly compared. Scale bar: 20 µm.

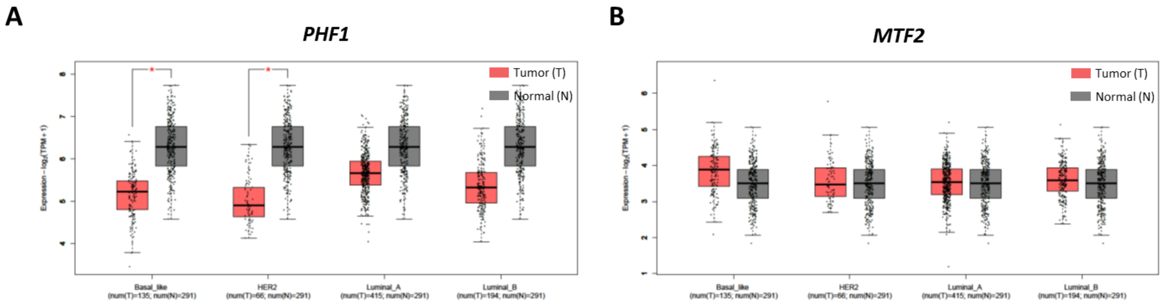


**Figure S3: PCL family gene expression analysis across a TCGA BRCA cohort sorted by breast cancer subtype.** Box plots display the expression levels of *PHF1* (**A**) and *MTF2* (**B**) in tumor (red) and normal (gray) samples for the indicated breast cancer subtypes (Basal-like, HER2, Luminal A, and Luminal B). Data are derived from TCGA/GTEx datasets and visualized using GEPIA2. Statistical significance between tumor and normal samples was determined by unpaired t-test (*$p < 0.05$). The number of tumor (T) and normal (N) samples included in the analysis is indicated below each plot. n= 291 (Normal), 194 (Luminal B), 415 (Luminal A), 66 (HER2), 135 (Basal-like).

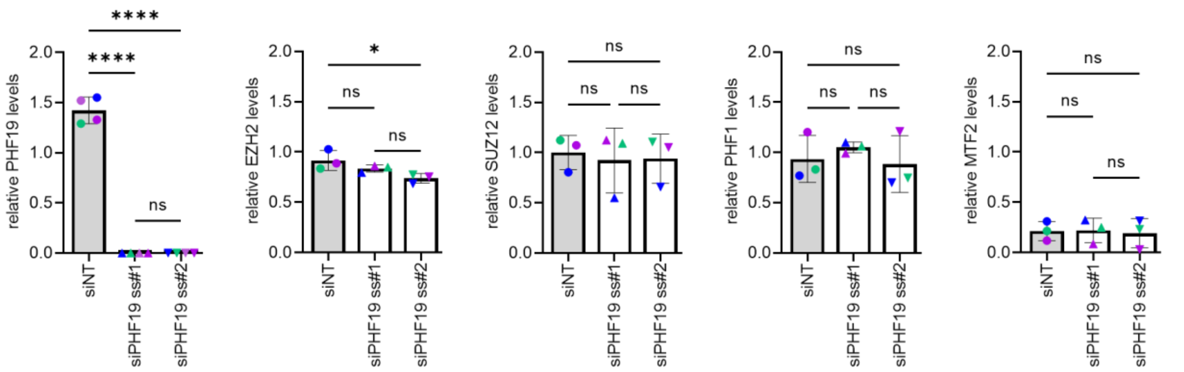


**Figure S4: Quantification of PRC2 subunit protein expression in MDA-MB-436 cells with PHF19 depletion.** MDA-MB-436 cells were transfected with the indicated siRNAs and lysed 96 hours later for Western blot analysis using the specified antibodies. GAPDH was used as a loading control. Bar diagrams represent the densitometric analysis of PHF19, EZH2, SUZ12, PHF1, and MTF2 protein levels in cell lysates obtained as described. Data are presented as measurements from n = 3 biological replicates, with biological repeats color-coded. Statistical significance was determined via one-way ANOVA testing: ****$p < 0.0001$; ns, not significant. Error bars indicate mean ± SD.

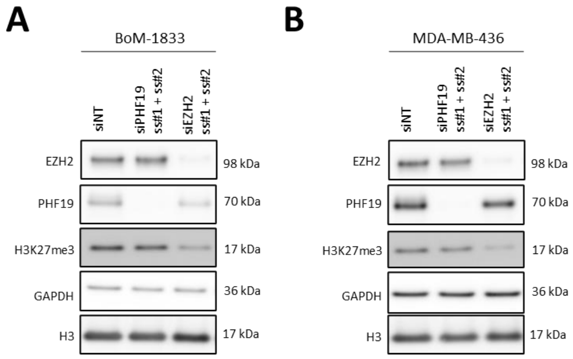


**Figure S5: EZH2 but not PHF19 depletion reduces global H3K27me3 levels in TNBC cells.** BoM-1833 (**A**) MDA-MB-436 (**B**) cells were transfected with the indicated siRNAs and lysed 96 hours post-transfection for Western blot analysis using the specified antibodies. GAPDH and H3 were used as loading controls.

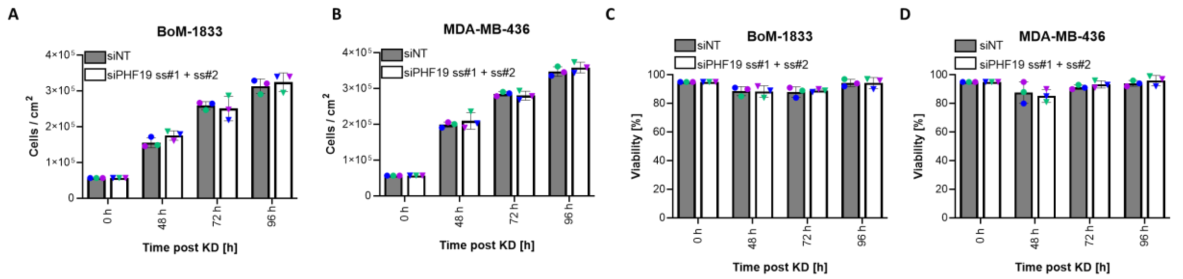


**Figure S6: PHF19 depletion does not affect cell growth and viability in TNBC cells.** BoM-1833 (**A, C**) and MDA-MB-436 (**B, D**) were transfected with the indicated siRNAs. The cells were harvested at the indicated time points after knockdown, counted (A, B) or subjected to viability measurements using trypan blue staining (C, D). Data are presented as measurements from n = 3 biological replicates, with biological repeats color-coded. Statistical significance was determined via two-way ANOVA testing, no significant differences within the matching timepoints were detected among siNT and siPHF19 knockdown cells. Error bars indicate mean ± SD.

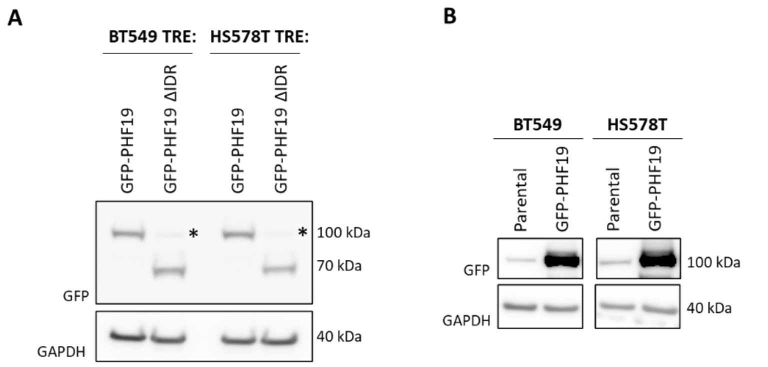


**Figure S7: Characterization of BT549 and HS578T cell models with inducible expression of the GFP-PHF19 variants.** (**A**) BT549 and HS578T cells with stable, inducible expression of GFP-PHF19 and GFP-PHF19 ΔIDR were treated with doxycycline for 24 hours, followed by harvesting and Western blot analysis of the lysates using the indicated antibodies. GAPDH was used as a loading control. An asterisk marks a signal observed at approximately 100 kDa, which was identified as unspecific antibody binding (see panel B for further details). (**B**) Western blot analysis comparing lysates from parental BT549 and HS578T cell lines with those of cell lines containing inducible GFP-PHF19. A GFP signal is observed in parental cells at approximately 100 kDa, indicating cross-reactivity of the GFP antibody with other cellular proteins running at the size of the GFP-PHF19 protein.

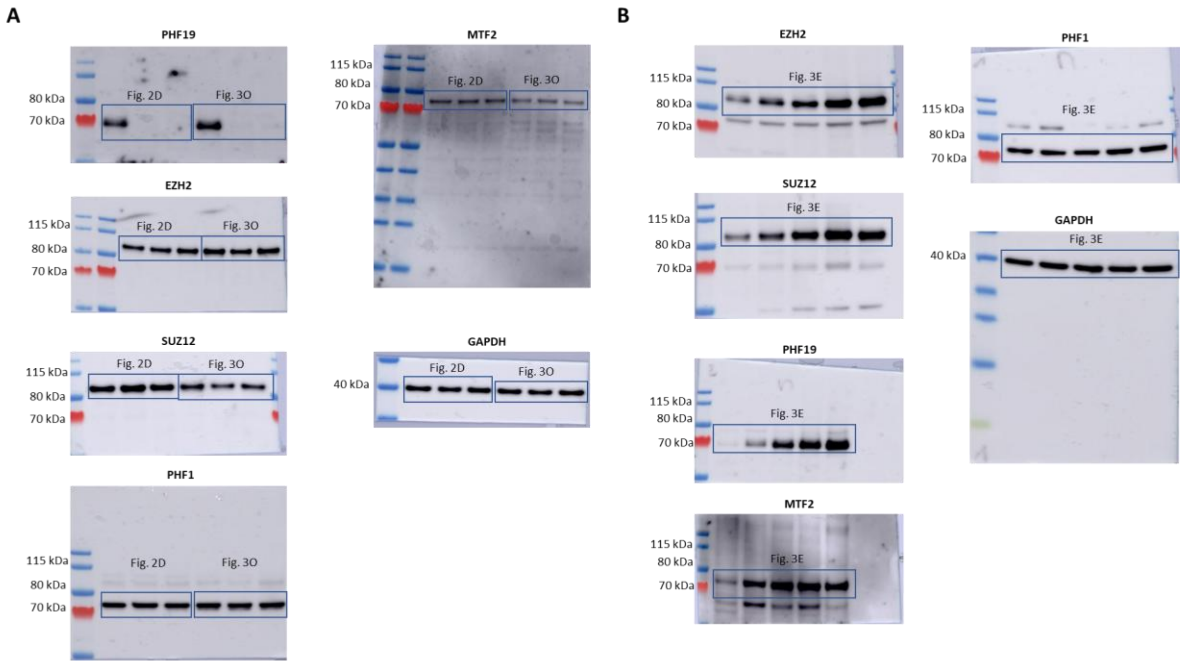


**Figure S8: Uncropped Western Blot images for Figures 2 and 3.** Uncropped Western blot images corresponding to the data presented in Figures 2 and 3. Merged JPEG images display the colorimetric molecular weight marker overlaid with the chemiluminescence signal. Rectangles indicate the regions of the membrane that were cropped in the TIFF files and shown in the annotated main figures. (**A**) Uncropped Western blots corresponding to Figures 2D and 3O. (**B**) Uncropped Western blots corresponding to Figure 3E.

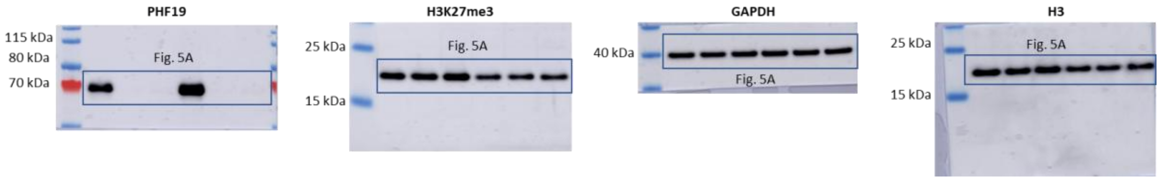


**Figure S9: Uncropped Western Blot Images for Figure 5A.** Merged JPEG images display the colorimetric molecular weight marker overlaid with the chemiluminescence signal. Rectangles indicate the regions of the membrane that were cropped in the TIFF files and shown Figure 5A.

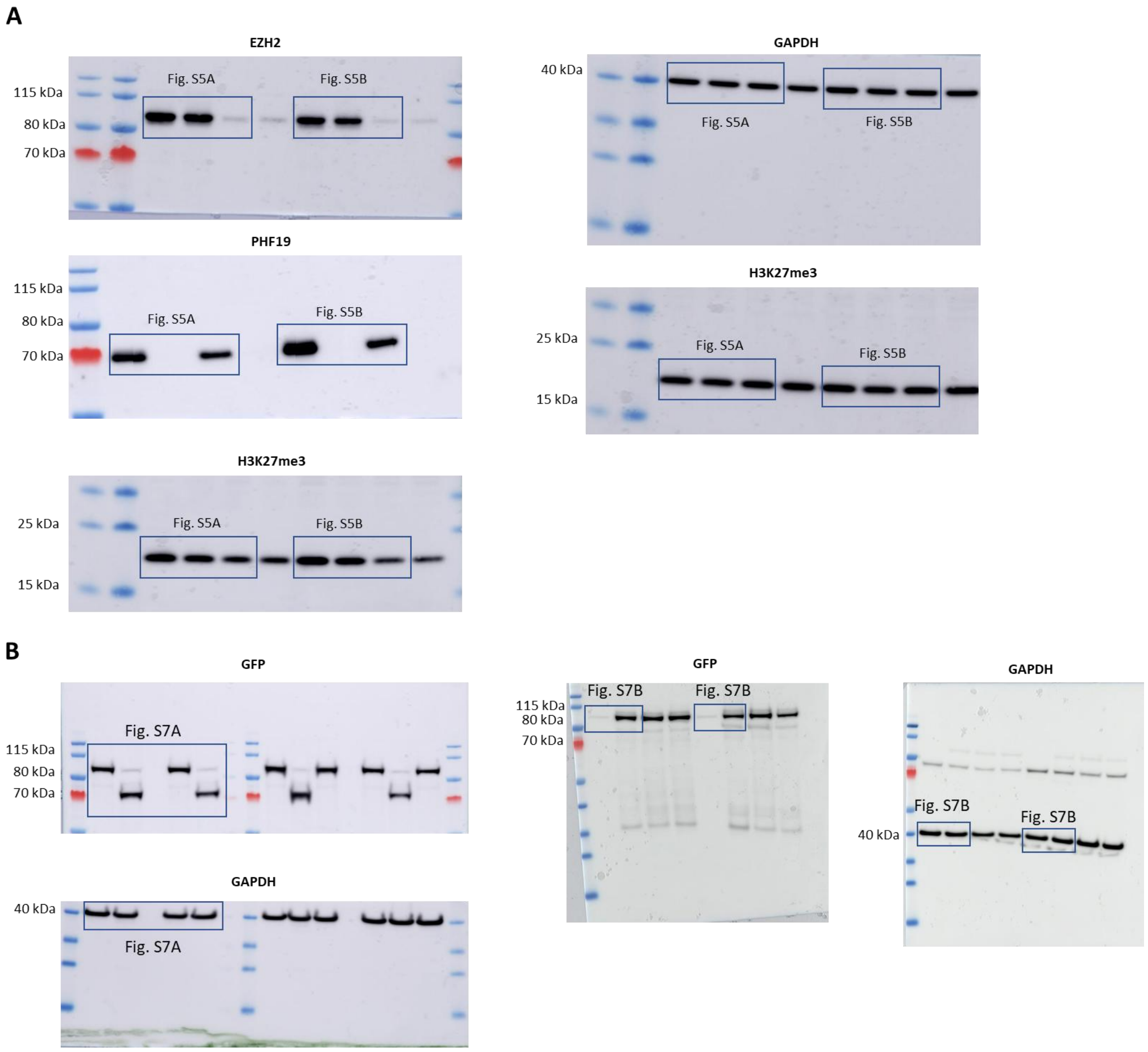


**Figure S10: Uncropped Western Blot images for Figures S5 and S7.** Uncropped Western blot images corresponding to the data presented in Figures S5 (**A**) and S7 (**B**). Merged JPEG images display the colorimetric molecular weight marker overlaid with the chemiluminescence signal. Rectangles indicate the regions of the membrane that were cropped in the TIFF files and shown in the annotated supplementary figures.
